# Inhibitors of the PqsR Quorum-Sensing Receptor Reveal Differential Roles for PqsE and RhlI in Control of Phenazine Production

**DOI:** 10.1101/2025.02.10.637488

**Authors:** Julie S. Valastyan, Emilee E. Shine, Robert A. Mook, Bonnie L. Bassler

## Abstract

*Pseudomonas aeruginosa* is a leading cause of hospital-acquired infections and it is resistant to many current antibiotic therapies, making development of new anti-microbial treatments imperative. The cell-to-cell communication process called quorum sensing controls *P. aeruginosa* pathogenicity. Quorum sensing relies on the production, release, and group-wide detection of extracellular signal molecules called autoinducers. Quorum sensing enables bacteria to synchronize group behaviors. *P. aeruginosa* possesses multiple quorum-sensing systems that control overlapping regulons, including those required for virulence and biofilm formation. Interventions that target *P. aeruginosa* quorum-sensing receptors are considered a fruitful avenue to pursue for new therapeutic advances. Here, we developed a *P. aeruginosa* strain that carries a bioluminescent reporter fused to a target promoter that is controlled by two *P. aeruginosa* quorum-sensing receptors. The receptors are PqsR, which binds and responds to the autoinducer called PQS (2-heptyl-3-hydroxy-4(1H)-quinolone) and RhlR, which binds and responds to the autoinducer called C4-HSL (C4-homoserine lactone). We used this reporter strain to screen >100,000 compounds with the aim of identifying inhibitors of either or both the PqsR and RhlR quorum-sensing receptors. We report results for 30 PqsR inhibitors from this screen. All of the identified compounds inhibit PqsR with IC_50_ values in the nanomolar to low micromolar range and they are readily docked into the autoinducer binding site of the PqsR crystal structure, suggesting they function competitively. The majority of hits identified are not structurally related to previously reported PqsR inhibitors. Recently, RhlR was shown to rely on the accessory protein PqsE for full function. Specifically, RhlR controls different subsets of genes depending on whether it is bound to PqsE or to C4-HSL, however, the consequences of differential regulation on the quorum-sensing output response have not been defined. PqsR regulates *pqsE*. That feature of the system enabled us to exploit our new set of PqsR inhibitors to show that RhlR requires PqsE to activate the biosynthetic genes for pyocyanin, a key *P. aeruginosa* virulence factor, while C4-HSL is dispensable. These results highlight the promise of inhibition of PqsR as a possible *P. aeruginosa* therapeutic to suppress production of factors under RhlR-PqsE control.

## Introduction

*Pseudomonas aeruginosa* is a Gram-negative bacterium that is ubiquitously present in the environment. *P. aeruginosa* is also a significant threat to human health, notably to burn victims, immunocompromised individuals, and people suffering from cystic fibrosis^1^. The CDC lists *P. aeruginosa* as a serious threat, as it accounted for over 30,000 infections and an estimated 2,700 deaths in hospitalized patients in 2017^2^. *P. aeruginosa* pathogenicity is tied to its ability to form surface-associated communities called biofilms, which, once present, are notoriously difficult to eradicate^3,4^. Additionally, *P. aeruginosa* is resistant to many current antibiotics^5,6^. For these reasons, development of new therapeutics to combat *P. aeruginosa* is crucial^7^.

Ongoing research endeavors seek to conceptualize and develop antimicrobials that function by mechanisms that prevent or delay resistance development by bacterial pathogens. One attractive possibility is therapeutics that target bacterial pathogenicity traits without affecting bacterial growth rate. The notion is that pathogens will not as readily evolve immunity to “behavior modification” treatments compared to resistance acquisition against bactericidal/bacteriostatic treatments because the former do not impose as strict a selection for resistance development as do the latter^8,9^. One behavior that is considered for such possible intervention is quorum sensing – the process of cell-cell communication that bacteria use to monitor cell density and orchestrate collective behaviors^10,11^. Quorum sensing depends on the production, release, and population-wide-detection of extracellular signal molecules, termed autoinducers, that accumulate with increasing population density. At low cell density, the concentration of autoinducers is below the level required for detection, and under this condition, bacteria behave as individuals^10,11^. As cell density increases, autoinducers accumulate to the threshold required for binding to their cognate receptors. Autoinducer-receptor complexes launch signal transduction cascades that drive changes in expression of genes underpinning group behaviors^10,11^. Importantly, in the case of *P. aeruginosa*, toxin production and biofilm formation, crucial disease determinants, are quorum-sensing-controlled traits that are enacted at high cell density^12^.

There are three major quorum-sensing systems in *P. aeruginosa*, each consisting of an autoinducer-receptor pair. All three receptors are transcription factors that, following binding to their cognate autoinducers, bind DNA, and drive quorum-sensing-controlled target genes. First, LasR binds the acyl-homoserine lactone (HSL) autoinducer 3O-C12-HSL, which is synthesized by LasI^13,14^. The LasR-3O-C12-HSL complex activates expression of a regulon of downstream genes that includes *lasA* and *lasB*, both of which encode elastase enzymes that are members of the suite of *P. aeruginosa* pathogenicity factors^14^. Second, RhlR binds to and is activated by a different HSL, C4-HSL, synthesized by RhlI^15^. The RhlR-C4-HSL complex activates production of genes specifying additional virulence factors, including toxic small molecules (e.g., phenazines and hydrogen cyanide), extracellular proteases, and rhamnolipids^16^. Finally, PqsR (also referred to as MvfR) binds to and is activated by the autoinducer 2-heptyl-3-hydroxy-4(1H)-quinolone, or *Pseudomonas* quinolone signal (PQS), as well as its precursor 2-heptyl-4(1*H*)-quinolone (HHQ)^17,18^. PQS is synthesized by enzymes encoded in the *pqsABCDE* operon, transcription of which is controlled by the PqsR-PQS complex^19^. Together, the PqsABCDE enzymes produce a family of quinolone molecules implicated in multiple *P. aeruginosa* roles including pathogenicity, iron chelation, and eukaryotic cell cytotoxicity^20^. Very importantly, PqsE is not required for synthesis of the PQS autoinducer^21^.

Beyond regulating virulence-mediating genes, all three *P. aeruginosa* quorum-sensing autoinducer-receptor pairs influence the expression and/or activity of the other quorum-sensing receptors. First, the LasR-3O-C12-HSL complex activates expression of *rhlR* and *pqsR*, encoding the other two quorum-sensing receptors^22,23^. This regulatory arrangement suggested that targeting LasR for inhibition could globally abrogate *P. aeruginosa* quorum sensing. Indeed, much work has focused on developing LasR inhibitors. However, during *P. aeruginosa* infection, *lasR* often acquires inactivating mutations, while quorum-sensing behavior is maintained through the Rhl system. Thus, inhibition of LasR does not currently appear to be a particularly promising strategy to control *P. aeruginosa* virulence^24–26^. Rather, efforts to inhibit *P. aeruginosa* quorum-sensing receptors are now focused on RhlR and PqsR as targets.

The PqsR-PQS complex promotes increased RhlR activity through induction of *pqsE* expression (Figure 1A)^27–30^. PqsE binds RhlR and facilitates its interaction with particular target promoters^31–34^. By contrast, RhlR suppresses PqsR activity by repressing expression of *pqsABCDE*, which reduces production of the PQS autoinducer (Figure 1A)^35,36^. Together, these mechanisms are proposed to affect the timing of induction of different quorum-sensing target genes. The absolute requirement for PqsE for expanded RhlR activity, coupled with the dependence of *pqsE* expression on the PqsR-PQS complex, makes PqsR an especially attractive target for small molecule inhibition.

**Figure 1.**
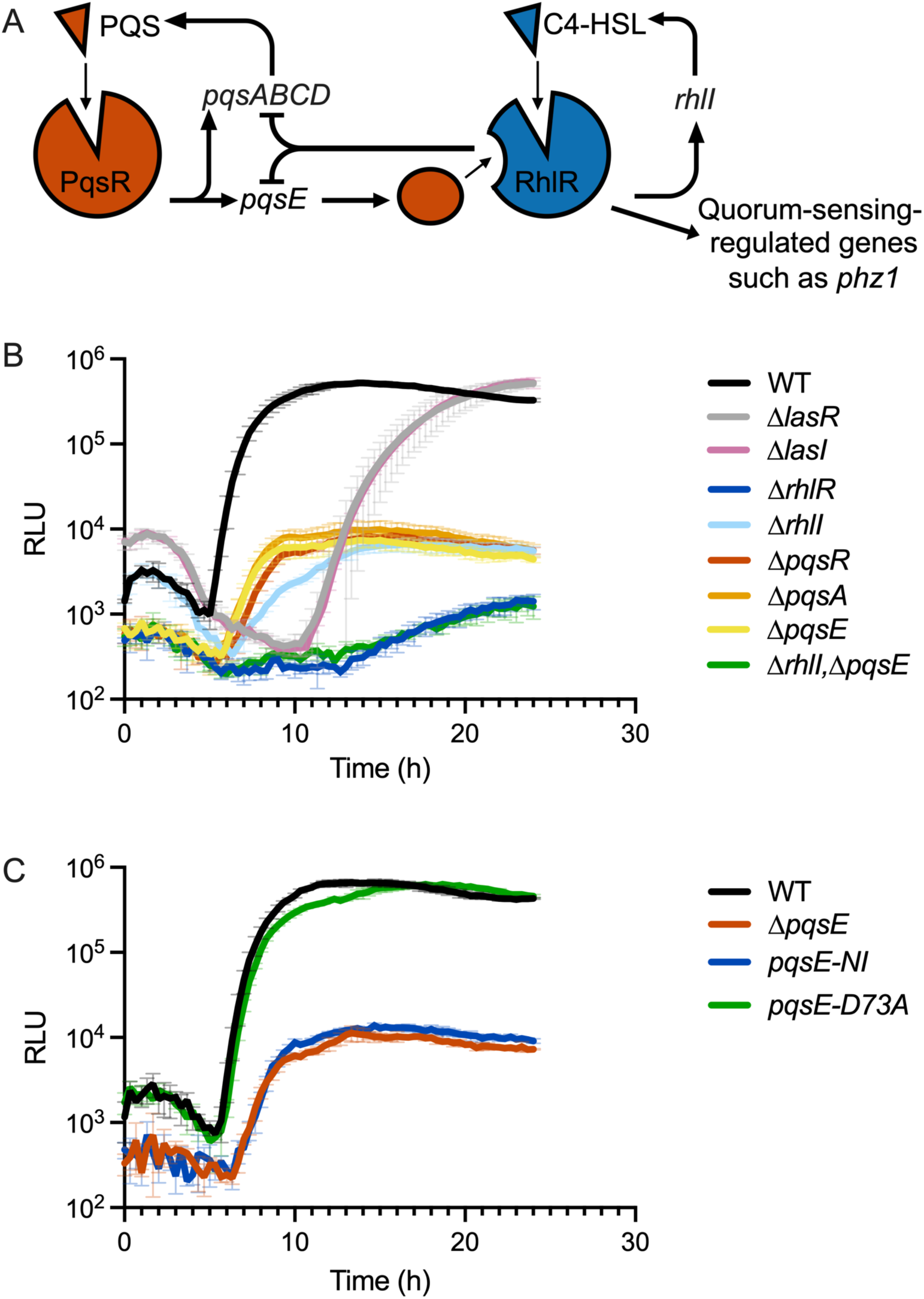
(A) Schematic summarizing PqsR- and RhlR-driven quorum sensing. (B, C) Light production over time from the p*phz1*-*lux* reporter in the designated *P. aeruginosa* strains. In B and C, light production is shown as relative light units (RLU), which is bioluminescence/OD_600_. Error bars = standard deviations of biological replicates, *n* = 3.

We constructed a bioluminescent transcriptional reporter fusion to a phenazine gene (p*phz1-lux*) that is regulated by both RhlR and PqsR and used it in a small molecule screen to identify inhibitors of RhlR and PqsR. Here, we report the results for the PqsR inhibitors. Characterization of the RhlR inhibitors will be reported elsewhere. Thirty PqsR inhibitors were uncovered in the screen. IC_50_ values were quantified in the above *P. aeruginosa* p*phz1-lux* reporter assay and in a recombinant *Escherichia coli* PqsR-specific reporter assay. Multiple compounds inhibited PqsR in the low micromolar range, with some compounds exhibiting nanomolar IC_50_ values. All tested compounds functioned competitively with PQS suggesting they bind in the ligand binding site. The 30 inhibitors span a surprisingly large chemical space. Docking analyses suggest how the diverse compounds can nonetheless bind in the same site. The compounds, by interfering with PqsR-PQS transcriptional activity, suppress PqsE production, but importantly they do not affect C4-HSL production. Therefore, the compounds only affect the PqsE-RhlR arm of the quorum-sensing circuit but not the RhlR-C4-HSL arm that functions independently of PqsE. This finding allowed us to show that *P. aeruginosa* employs PqsE-RhlR and RhlR-C4-HSL to differentially control production of phenazine compounds, some of which are key *P. aeruginosa* virulence factors.

## Results

### Establishing chromosomal bioluminescent transcriptional fusions that report on quorum-sensing-controlled genes in P. aeruginosa

To maximize the probability of uncovering small molecule inhibitors of the different *P. aeruginosa* quorum-sensing pathways, we relied on published RNAseq analyses to guide identification of genes controlled by multiple *P. aeruginosa* quorum-sensing pathways. Based on the data^16^, we constructed luciferase fusions to promoters driving five quorum-sensing-regulated genes, *chiC*, *hcnA*, *rhlA*, *phz1*, and *phz2*. We also made fusions to two promoters driving non-quorum-sensing-regulated genes *rpsL* and *tac*, as controls. To assess the dynamic ranges of these *lux* reporters, and their reliance on the different *P. aeruginosa* quorum-sensing systems, we monitored time courses of luciferase production in wildtype (WT) *P. aeruginosa* PA14 and in *P. aeruginosa* PA14 strains lacking a single quorum-sensing receptor (τ1*lasR*, τ1*rhlR*, or τ1*pqsR*) (Figures 1B and S1). As expected, no quorum-sensing-driven regulation occurred for p*rpsL-lux* and p*tac-lux* in any strain (Figure S1), making them excellent control fusions that could additionally be used to identify and eliminate compounds that inhibit luciferase from our screen. By contrast, the p*chiC-*, p*hcnA-*, p*rhlA-*, p*phz1-*, and p*phz2-lux* promoter fusions all showed significant reductions in light output in each of the *P. aeruginosa* quorum-sensing deficient strains (the p*phz1-lux* data are shown in Figure 1B, data for the other reporters are provided in Figure S1).

Due to its superior dynamic range in response to regulation by RhlR and PqsR, we chose the p*phz1-lux* reporter to use in our small molecule inhibitor screen. To expand our examination of its responses to quorum-sensing perturbation, we engineered additional mutations into the p*phz1-lux* reporter strain. The phenotypes of mutants deleted for the *lasI* or *pqsA* synthase gene mimicked those of the strains deleted for the cognate receptor gene, *lasR* or *pqsR,* respectively. By contrast, p*phz1-lux* expression in the τ1*rhlR* mutant differed from that in the τ1*rhlI* mutant (Figure 1B). This difference occurs because both the RhlI-produced autoinducer, C4-HSL, and PqsE contribute to RhlR activity and consequently, RhlR differentially regulates p*phz1* depending on whether it is bound to C4-HSL, to PqsE, or to both C4-HSL and PqsE^16,27^. Supporting this assertion are our data showing that while p*phz1-lux* expression in the τ1*pqsE* and Δ*rhlI* mutants differs from that in the τ1r*hlR* mutant, p*phz1-lux* expression in the double τ1*rhlI* τ1*pqsE* mutant is indistinguishable from that in the τ1*rhlR* mutant (Figure 1B). These data verify that both RhlI and PqsE are required for maximal activation of p*phz1* by RhlR.

PqsE acts as both a thioesterase and an accessory protein^31,33^. The PqsE accessory function is protein-protein interaction with RhlR and interaction is required for RhlR to activate a subset of *P. aeruginosa* virulence genes, including *phz1*. The PqsE thioesterase activity is dispensable for this accessory function^31^. PqsE mutants exist that possess one, the other, or neither the thioesterase and/or protein-protein interaction functions, showing that the activities are separable. Germane to this work are two PqsE mutants: First, a PqsE mutant containing alanine substitutions in place of three arginine residues that render PqsE unable to bind to RhlR^33^. This PqsE mutant is called PqsE-NI (NI for non-interacting). PqsE-NI is unable to enhance RhlR-driven gene expression while its thioesterase activity remains intact. Conversely, the PqsE-D73A mutant, which harbors an alteration in an essential active site catalytic residue, possesses no thioesterase activity but retains the ability to interact with and enhance RhlR-mediated gene control^31^. Figure 1C shows that PqsE-NI does not enhance RhlR activation of p*phz1-lux* while PqsE-D73A increases RhlR-directed p*phz1-lux* activation to the level of WT PqsE. These results confirm that it is the PqsE-RhlR interaction and not the PqsE enzymatic activity that is crucial for regulation of p*phz1-lux*.

### A screen for small molecule inhibitors of RhlR- and PqsR-directed quorum sensing

Suppressing traits in *P. aeruginosa* using small molecules has proven challenging because *P. aeruginosa* possesses mechanisms that prevent the uptake of compounds and efflux pumps that export molecules once internalized^37^. To circumvent issues with non-uptake/efflux, we performed our inhibitor screen in the presence of an additive called SPR741^38^, a polymyxin B derivative that partially permeabilizes the bacterial outer membrane, thereby promoting enhanced entry of small molecules. Our rationale for using SPR741 in the initial screen was to maximally reveal compounds of potential interest that would otherwise be undetectable due to impermeability. We reasoned that if needed, further refinements could be made to interesting compounds to enhance penetration in the absence of SPR741.

To establish optimal conditions for the small molecule inhibitor screen, we exploited a compound that we uncovered earlier that, via an uncharacterized mechanism, inhibits luciferase. We call this compound 368A. Figure 2A shows the *P. aeruginosa* p*phz1-lux* output following administration of 100 µM 368A in the presence and absence of different concentrations of SPR741. Inclusion of SPR741 at 8 µg/mL increases 368A inhibitory potency by 5-fold. To confirm that 8 µg/mL SPR741 did not significantly alter quorum sensing in *P. aeruginosa*, we tested its effects on our set of quorum-sensing mutants containing the p*phz1-lux* reporter. In all cases, the same patterns of p*phz1-lux* expression occur with and without SPR741 (compare data in Figure 2B to that in Figure 1B). Below, all screening steps were performed in the presence of 8 µg/mL SPR741.

**Figure 2.**
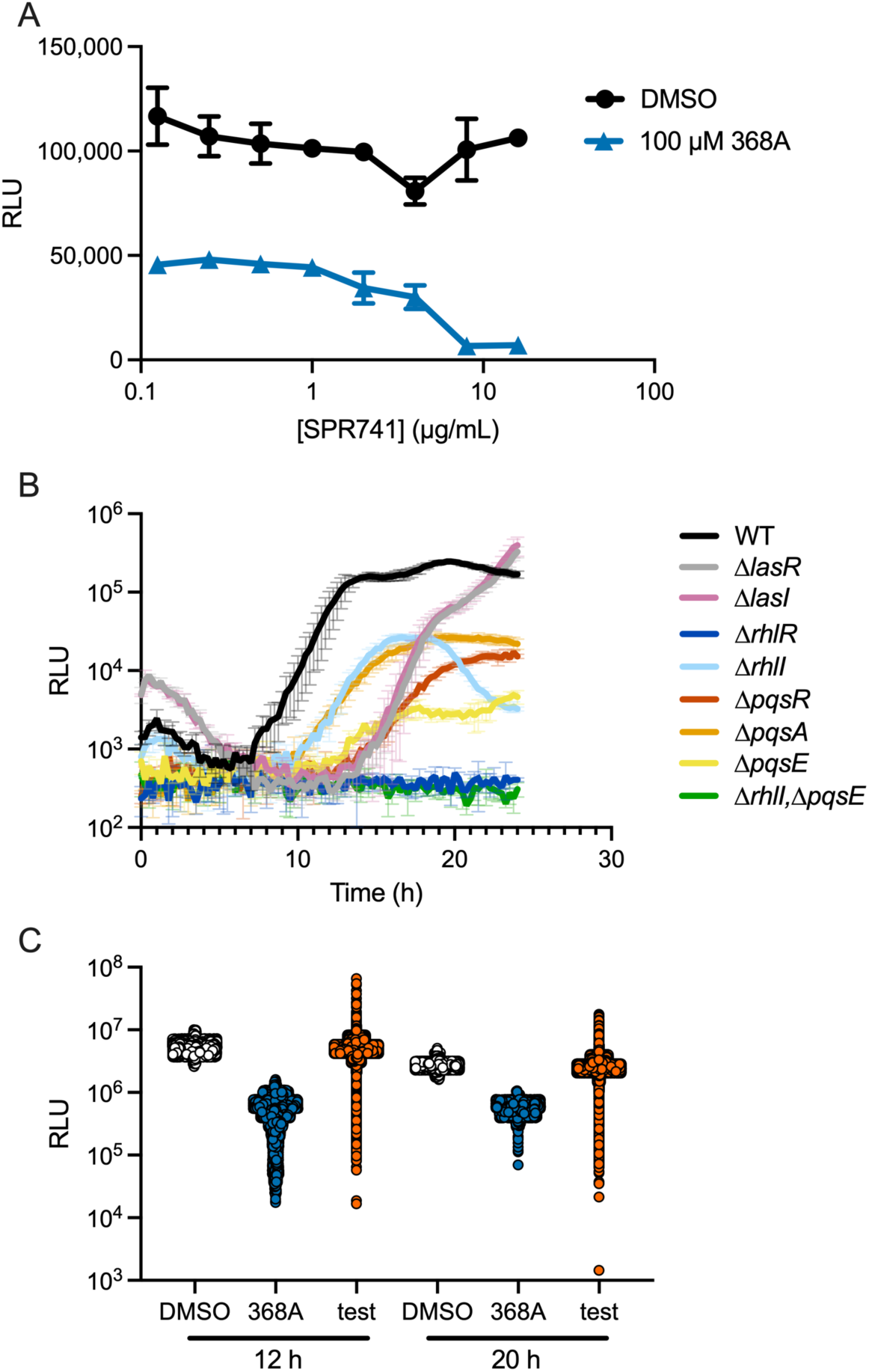
(A) Light production from WT *P. aeruginosa* carrying the p*phz1*-*lux* reporter at the designated SPR741 concentrations following 10 h of growth. (B) Light production from the p*phz1*-*lux* reporter over time in the designated *P. aeruginosa* strains in the presence of 8 µg/mL SPR741. (C) Light production from WT *P. aeruginosa* carrying the p*phz1*-*lux* reporter in the presence of DMSO (white, *n* = 5,200), 40 µM 368A (blue, *n* = 5,200), or the screening compounds (orange, *n* = 104,000), excluding control outliers based on the WuXi AppTec quality control algorithm. RLU as in Figure 1. In panels A and B, error bars = standard deviations of biological replicates, *n* = 3.

### *A screen of 100,000 small molecules for* P. aeruginosa *RhlR and PqsR inhibitors*

A small molecule screen was performed at WuXi AppTec in 384-well format as follows: Sixteen wells in each plate were administered DMSO as negative controls and 16 wells were administered 40 µM of the 368A luciferase inhibitor as positive controls. Test compounds (final concentration 40 µM and 0.4% v/v) were administered to the other wells. Liquid LB growth medium containing 8 µg/mL SPR741 was supplied together with an aliquot of an overnight culture of WT *P. aeruginosa* carrying the p*phz1-lux* reporter that had been diluted to a final OD_600_ of 0.001.

Figure 2C shows that the output ranges from the negative and positive controls (white and blue, respectively) were maintained throughout the screen and that some of the over 100,000 test molecules (orange) inhibited p*phz1-lux* more potently than did the positive control 368A inhibitor (blue). Light production and OD_600_ (i.e., cell growth) were quantified at 12 and 20 h. Compound degradation and export often confound screening in *P. aeruginosa*. Thus, the 20 h timepoint allowed us to focus on molecules that exerted sustained effects. Assessing OD_600_ allowed us to eliminate compounds that retarded or arrested cell growth.

One thousand compounds from the screen were selected for retesting based on potency of p*phz1-lux* inhibition and diversity of compound structure (Figure 3A). Roughly 75% of the retested compounds showed comparable results to those obtained in the original screen. This subset of compounds was subsequently assessed for activity against the control p*tac-lux* fusion to eliminate luciferase inhibitors (Figure S2A). As expected, many compounds failed to move forward from this round. 22% (167 compounds) of the retested compounds inhibited p*phz1-lux* but did not significantly inhibit p*tac-lux*. These 167 remaining compounds were next examined by serial dilution to determine their IC_50_ values against p*phz1-lux*. From these analyses, 49 compounds were purchased for further in-house analyses.

**Figure 3.**
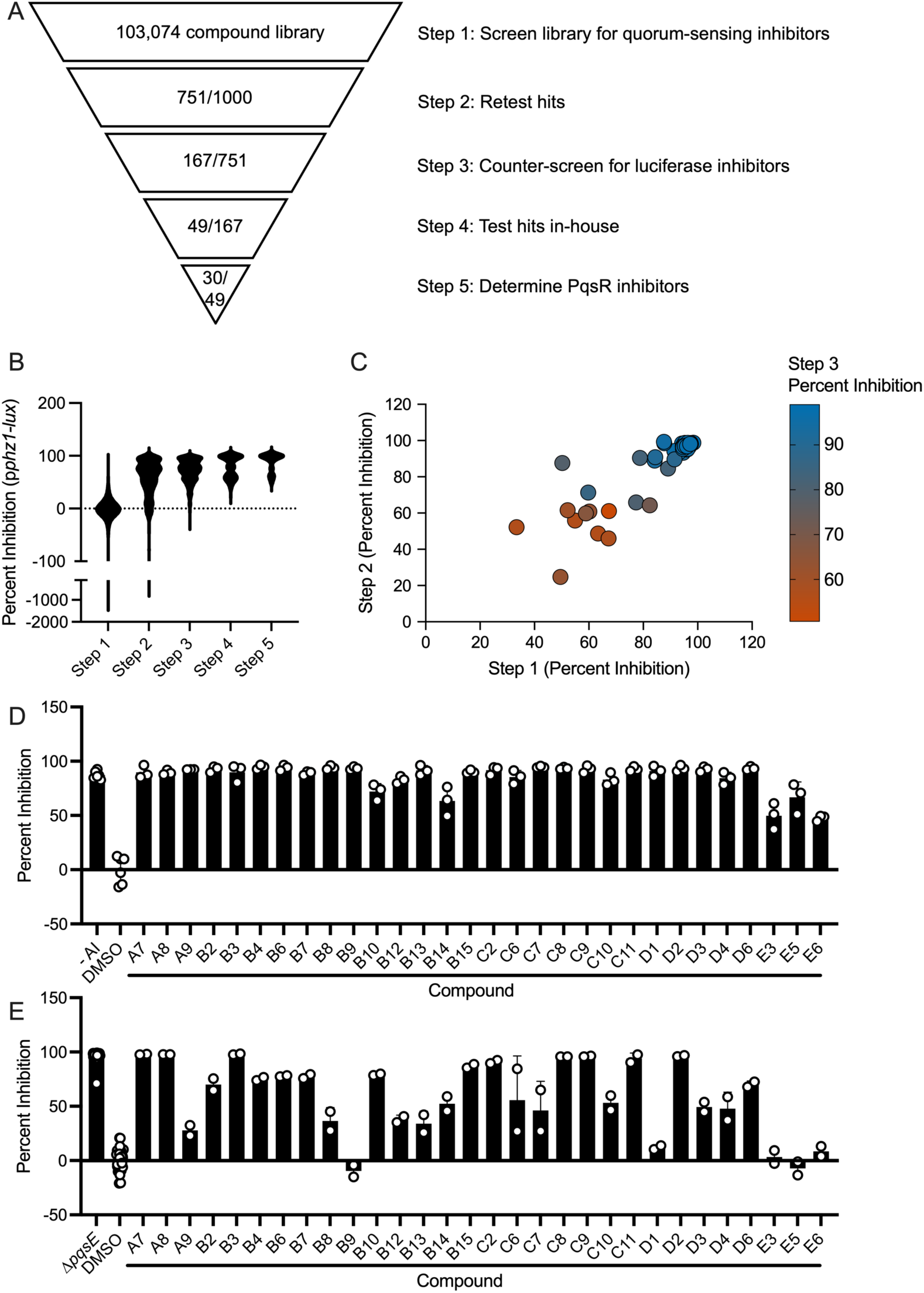
(A) Summary of results from each step of the small molecule screen for inhibitors of the p*phz1*-*lux* reporter in WT *P. aeruginosa*. (B) Inhibition of light production from the p*phz1*-*lux* reporter in WT *P. aeruginosa* by all compounds remaining at each step of the small molecule screen. (C) Inhibition of light production from the p*phz1*-*lux* reporter in WT *P. aeruginosa* by the final 49 candidate compounds in three different steps of the screen. (D) Inhibition of light production from the p*pqsA*-*lux* reporter by 100 μM of the designated candidate compounds in *E. coli*. The left-most bar shows the result when no PQS autoinducer was added. All other bars show the results when PQS was present at 100 nM. Note SPR741 was not included in this assay. (E) Inhibition of pyocyanin production in WT *P. aeruginosa* by 100 µM of the designated candidate compounds in the presence of SPR741. The left-most bar shows the result for the τ1*pqsE P. aeruginosa* mutant as the positive control and the second bar shows the result for DMSO as the negative control. In panels D and E, error bars = standard deviations of biological replicates, *n* = 3.

To assess overall success of the screening pipeline, we quantified the average potency of all compounds remaining at each successive step as well as the reproducibility of the activities of the final 49 selected compounds. Following each screening step, the average percent inhibition of p*phz1-lux* increased, suggesting that our funneling criteria delivered increasingly potent molecules (Figure 3B). Additionally, the 49 hit compounds showed strong reproducibility at each step (Figure 3C) and they did not significantly inhibit luciferase in the p*tac-lux* control screen (Figure S2B). Together, these data make the 49 selected compounds promising leads as specific *P. aeruginosa* quorum-sensing inhibitors.

### Identification of PqsR as the target of the small molecule inhibitors

To determine the target(s) of the 49 hit compounds in *P. aeruginosa,* we assayed them in a set of *E. coli*-based reporter strains that read out exclusively RhlR, LasR, or PqsR function^39^. The three assays work by a similar logic. In each case, the gene encoding one of the three quorum-sensing receptors is driven by an arabinose-inducible promoter. The receptor proteins, once made, are activated by exogenously supplied cognate autoinducer. The receptor-autoinducer complexes activate a target gene promoter fused to luciferase (p*rhlA-lux* for RhlR, p*lasB-lux* for LasR, and p*pqsA-lux* for PqsR). Surprisingly, at 100 µM, 30 of the 49 compounds potently inhibited PqsR (Figure 3D). Some of these compounds also showed modest inhibition of RhlR (Figure S3A) and LasR (Figure S3B). PqsR has previously been pinpointed as a target for quorum-sensing inhibition. Our findings reinforce the hypothesis that PqsR is surprisingly amenable to inhibition in *P. aeruginosa*, especially compared to RhlR, which has proven surprisingly difficult to inhibit with small molecules. Below, we focus on these 30 compounds that target PqsR. The other 19 compounds will be characterized at a later time.

Pyocyanin is a quorum-sensing-controlled blue-green-colored toxic compound that *P. aeruginosa* produces at high cell density^40,41^. Its production is activated by PqsR-PQS^42,43^. Thus, reduced expression of the pyocyanin biosynthetic genes occurs in a τ1*pqsR* strain. Its striking color makes pyocyanin a convenient visible output of PqsR-PQS-driven quorum-sensing activation. We tested our panel of 30 PqsR inhibitors for the ability to reduce *P. aeruginosa* pyocyanin production. The τ1*pqsE* strain, which makes no pyocyanin, served as the positive control. Twenty-five of the 30 compounds inhibited pyocyanin production by at least 25% compared to when DMSO was added (Figure 3E). The pyocyanin assay requires 16 h of bacterial growth, so this result also indicates that the majority of the compounds must not be degraded or exported by *P. aeruginosa* since they displayed sustained PqsR inhibition. All of the compounds that inhibited pyocyanin production in the presence of SPR741 (Figure 3E) also showed at least 25% inhibition in its absence (Figure S4), suggesting that *P. aeruginosa* readily internalizes these molecules. A subset of the molecules, such as the compound denoted D1, showed improved pyocyanin inhibition in the absence of SPR741 compared to in its presence. Further experiments are required to understand the mechanism underlying this peculiar result.

Table 1 presents the structures of and IC_50_ values for the PqsR inhibitors identified here. The IC_50_ values were calculated from *P. aeruginosa* p*phz1-lux* and *E. coli* PqsR/p*pqsA-lux* assays. Notably, these compounds have diverse structures (Figure S5) showing that multiple chemotypes were identified with the capacity to inhibit PqsR. By far, hits containing amides represent the most frequent structural features, with secondary amides accounting for 18 and tertiary amides accounting for 8 of the 30 hits. Some hits identified here contain structural fragments resembling those in previously reported PqsR inhibitors^44–50^.

**Table 1.**
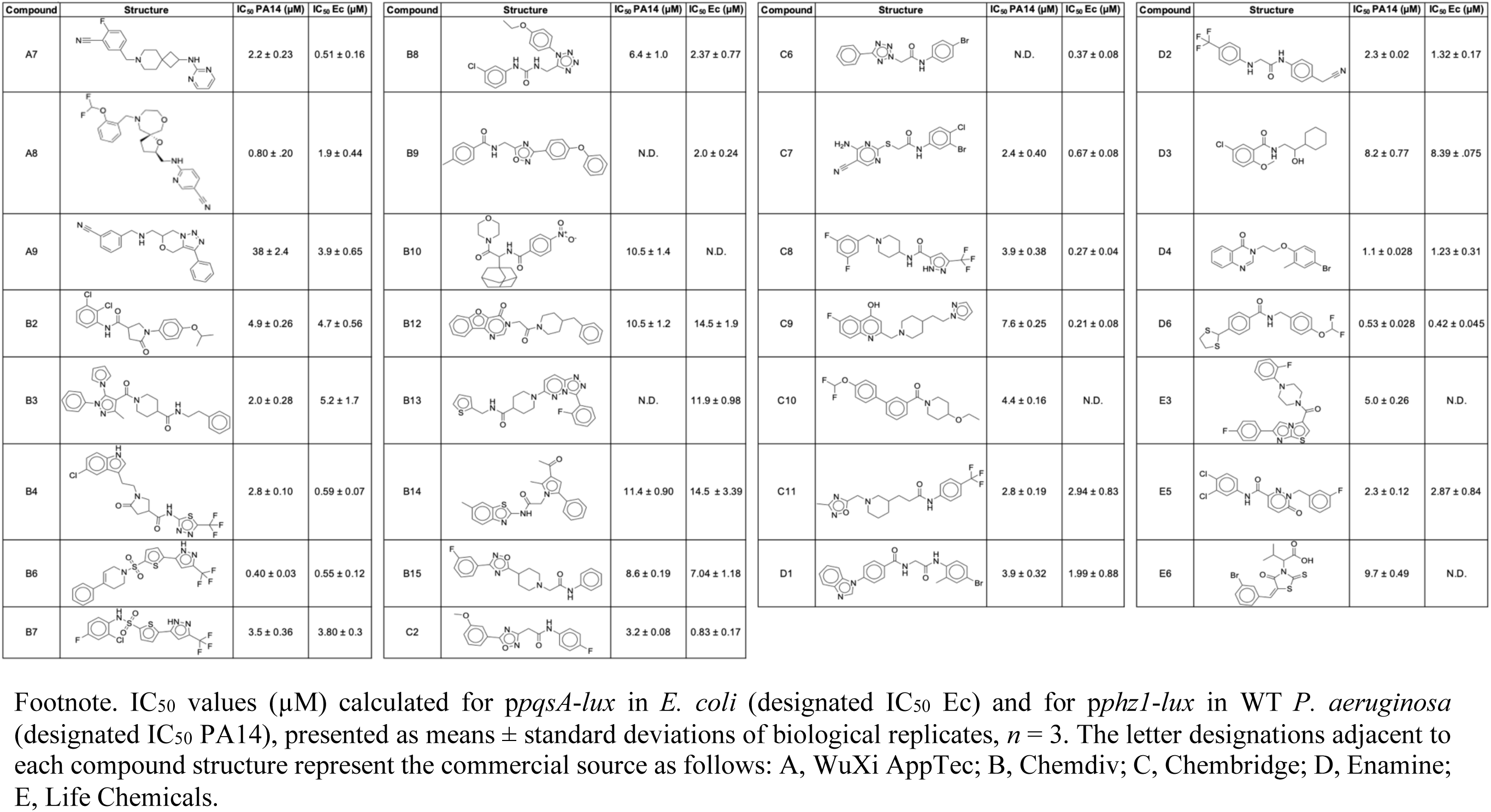
PqsR inhibitors uncovered in a small molecule screen.

There are at least 20 structures of PqsR in the PDB, many with inhibitors bound in the ligand binding pocket (Figure 4A, left). Aligning these structures shows that most of the different compounds fill much of the available space in the binding pocket (Figure 4A, middle). Ninety-degree rotation reveals that all of the compounds bind in a similar, overall L-shaped pose (Figure 4A, right)^46–49,51–56^. Using docking analyses, we examined whether our compounds likewise have the potential to bind in the ligand binding pocket^57–60^. We selected the structure of the native ligand HHQ bound to PqsR reported by Zender et al. as our reference (blue in Figure 4A,B)^47^. Docking the native HHQ ligand faithfully reproduced the binding pose and ligand conformation of HHQ observed in the x-ray structure, with a GScore^61^ of -6.9 (Figure 4B, left and Figure S6A). All of our compounds easily docked into the PqsR ligand binding site with GScores < -5.5 (Figure 4B, left, Figure S6, and Table S1), also filling the space (Figure 4B, middle) and in the L-shaped configuration (Figure 4B, right). Indeed, half of our compounds had GScores superior to that of HHQ (Table S1).

**Figure 4.**
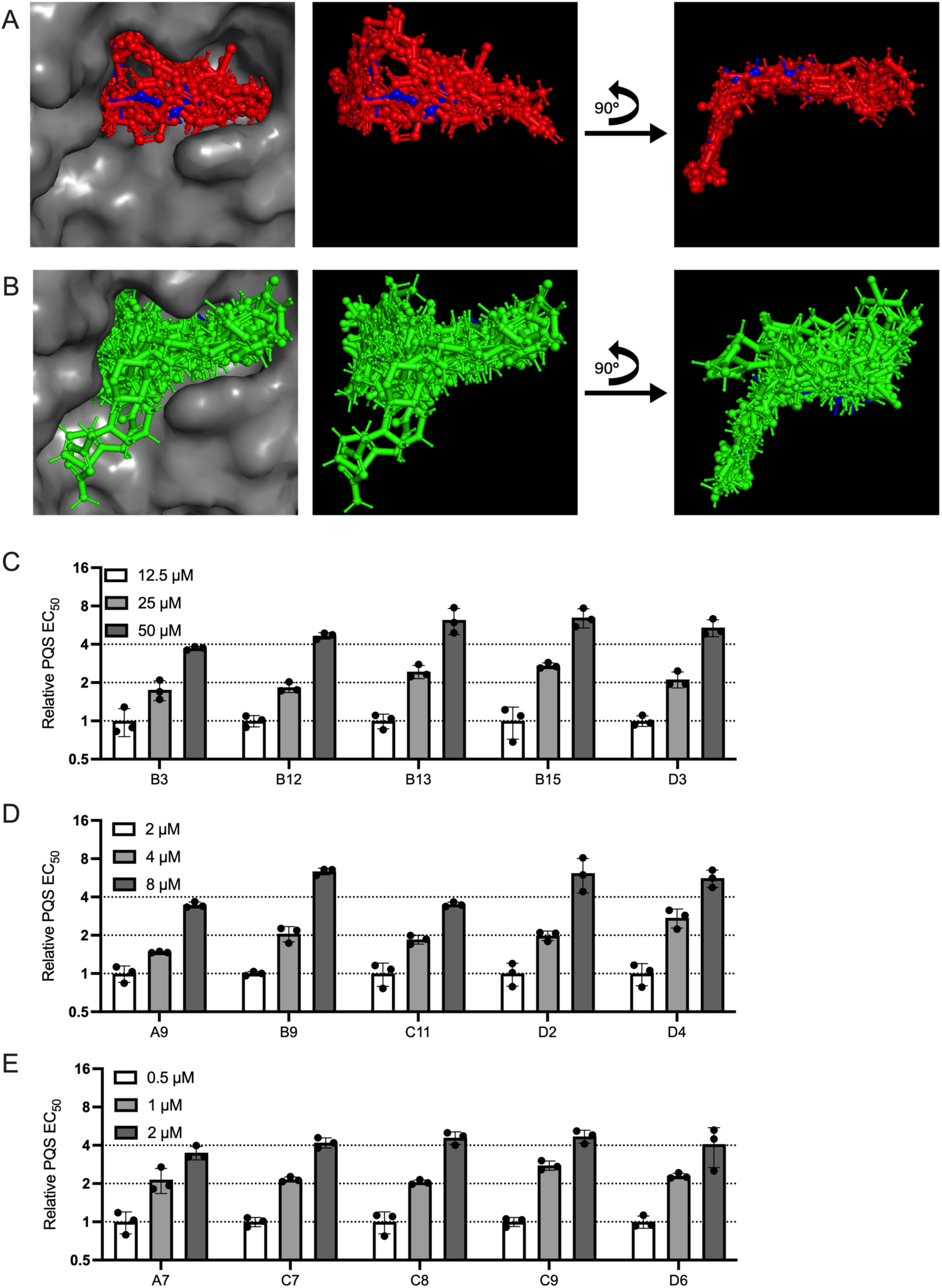
(A) Alignment of crystal structures of PqsR bound to small molecule agonists and antagonists. PDB files for aligned crystals are 4JVI^51^, 7O2T^62^, 7O2U^63^, 6TPR^46^, 6Z17^48^, 6Z5K^48^, 7QA0^49^, 7QA3^49^, 7QAV^49^, 6Z07^48^, 6Q7U^47^, 7NBW^48^, 6B8A^52^, 6Q7W^47^, 7P4U^55^, 8Q5L^56^, 6YIZ^53^, 8Q5K^56^, 6YZ3^48^, and 4JVD^51^. Gray = protein from 6Q7U, blue = HHQ from 6Q7U, red = all other small molecules. (B) Docking of our small molecule antagonists into PqsR crystal structure 6Q7U. Gray = protein from Q67U, blue = HHQ from 6Q7U, green = our small molecules. (C-E) Shown are relative EC_50_ values for PQS binding to PqsR in the presence of increasing concentrations of the specified PqsR inhibitors. Assays were performed with a subset of inhibitors that exhibited IC_50_ values (C) above 5 μM (D) between 1 and 5 μM, (E) less than 1 μM. In panels C-E, concentrations of inhibitors used are provided in the legends on the top left of each graph, guide lines show relative EC_50_ values of 1, 2, and 4 to aid in visualization, error bars = standard deviations of biological replicates, *n* = 3.

If the PqsR inhibitors identified here indeed bind in the ligand binding site, they should act competitively. To test this assumption, we assayed 15 of our compounds for competition with the PQS ligand for binding to PqsR using our *E. coli* PqsR reporter assay. EC_50_ values for PQS are provided for each concentration of inhibitor tested (Figure S6L). To visualize the fold changes between inhibitor concentrations, EC_50_ values relative to the lowest concentration of inhibitor are shown in Figure 4C-E. In every case, increasing the amount of inhibitor two-fold caused a corresponding two-fold increase in the EC_50_ for PQS. These data corroborate the docking studies and imply that our compounds are competitive inhibitors.

### In vivo small molecule PqsR quorum-sensing inhibition occurs by suppression of pqsE expression

The primary quorum-sensing role of PqsR-PQS in *P. aeruginosa* is to activate expression of the *pqsABCDE* operon, which includes *pqsE*^64^. As described above, PqsE enhances RhlR-DNA-binding activity, thereby assisting RhlR in carrying out a subset of its regulatory functions. In agreement with this mechanism, and as shown in Figures 1B and 5A, deletion of *pqsR* from *P. aeruginosa* severely diminishes p*phz1-lux* output. Expression of *pqsE* from a constitutive promoter (*pqsE-ON*) restores light production to the τ1*pqsR* mutant (Figure 5A). Thus, synthetically providing the PqsE needed to interact with RhlR to activate the p*phz1-lux* promoter overrides the need for PQS. We used this feature of the system to assess if a subset of our identified compounds affects *P. aeruginosa* quorum-sensing components in addition to inhibition of PqsR. To do this, we treated the *P. aeruginosa* Δ*pqsR pqsE-ON* strain carrying p*phz1-lux* with candidate compounds at 25 µM. The rationale is that any compounds capable of inhibiting p*phz1-lux* expression in this assay arrangement must affect a component other than PqsR since PqsR is absent. All assayed compounds lost the ability to suppress p*phz1-lux* expression when *pqsE* was constitutively supplied (compare data in Figure 5B to that in 5C) indicating that PqsR is the target affected by our compounds.

**Figure 5.**
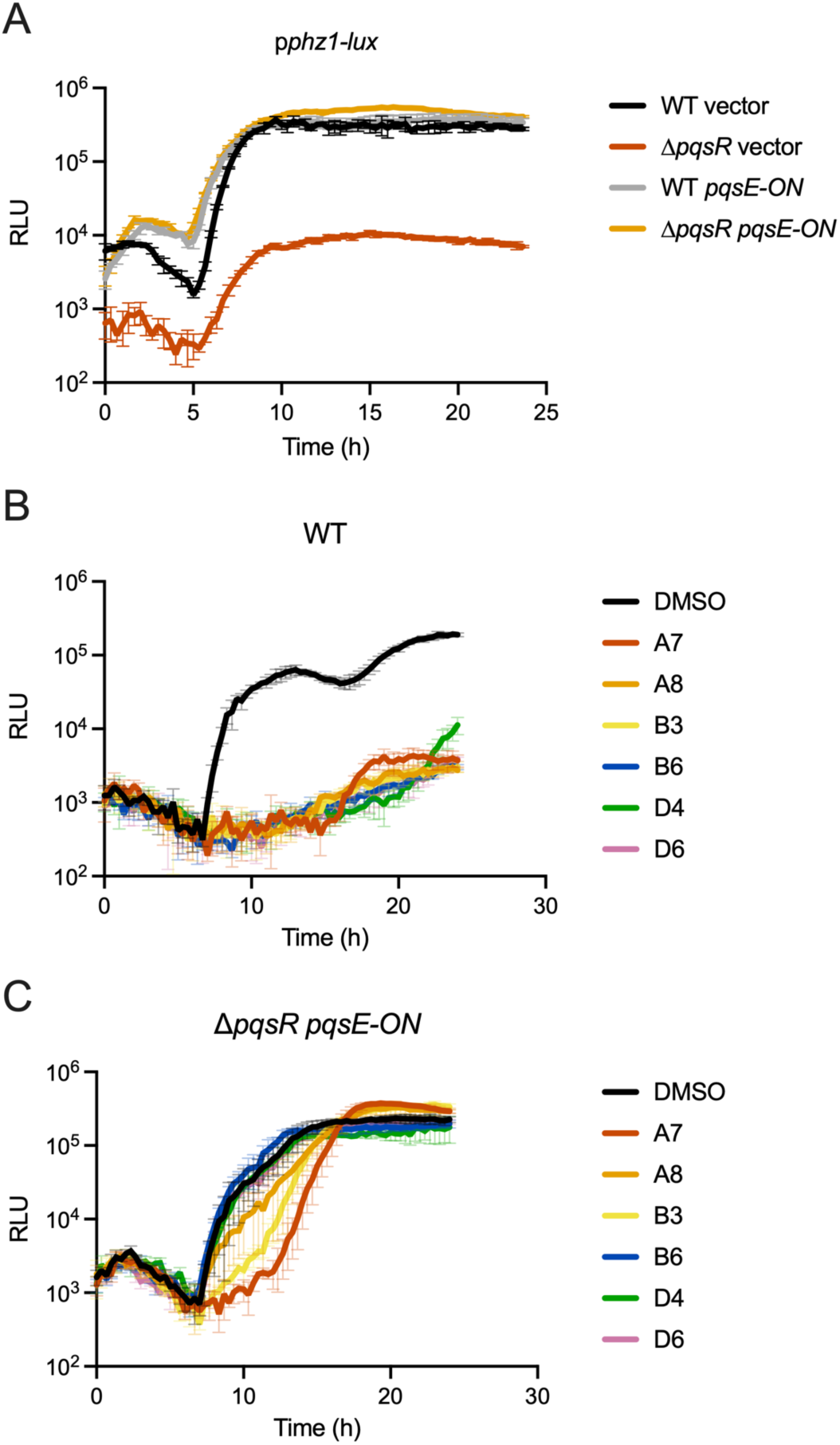
(A) Light production over time from the p*phz1*-*lux* reporter in the designated *P. aeruginosa* strains. The *pqsE-ON* construct constitutively produces PqsE. (B) Light production over time from the p*phz1*-*lux* reporter in WT *P. aeruginosa* in the presence of DMSO or the designated PqsR inhibitors. (C) Light production over time from the p*phz1*-*lux* reporter in the τ1*pqsR P. aeruginosa* strain constitutively producing *pqsE* in the presence of DMSO or the designated PqsR inhibitors. RLU as in Figure 1. In all panels, error bars = standard deviations of biological replicates, *n* = 3.

### Different combinations of phenazines are produced depending on whether RhlR is bound to C4-HSL or to PqsE

RhlR controls different subsets of genes depending on whether it is bound to C4-HSL or to PqsE^16,27^. One set of genes subject to differential RhlR regulation are the phenazine biosynthetic genes^16^. Synthesis of phenazines originates with the protein products of the *phz1* and *phz2* gene clusters^65^. Both operons produce enzymes that convert chorismic acid into phenazine-1-carboxylic acid (PCA)^65^. As shown throughout this work, RhlR partners with both C4-HSL and PqsE to control expression of the *phz1* operon. Analogous dual regulation occurs for the *phz2* operon. Distinct regulation by RhlR-C4-HSL and by PqsE-RhlR can be exemplified through examination of the patterns of regulation of two phenazine biosynthesis genes, *phzH* and *phzS* (Figure 6A)^16,27^. Specifically, *phzH* encodes a glutamine amidotransferase that converts PCA into phenazine-1-carboxamide (PCN)^65^. *phzH* expression depends more highly on RhlR-C4-HSL than on PqsE-RhlR^27^. By contrast, *phzS*, encoding the PhzS monooxygenase, is involved in converting PCA to pyocyanin and 1-hydroxyphenazine (1-OH-Phz)^65^. *phzS* is downregulated in the Δ*rhlR* mutant but not in the Δ*rhlI* mutant^16^, suggesting *phzS* control is more reliant on PqsE-RhlR than on RhlR-C4-HSL. However, we note that the role of PqsE in *phzS* regulation has not been directly examined. These data suggest that RhlR can drive production of different phenazines depending on the relative levels of C4-HSL and PqsE in the cell and/or to which it is bound.

**Figure 6.**
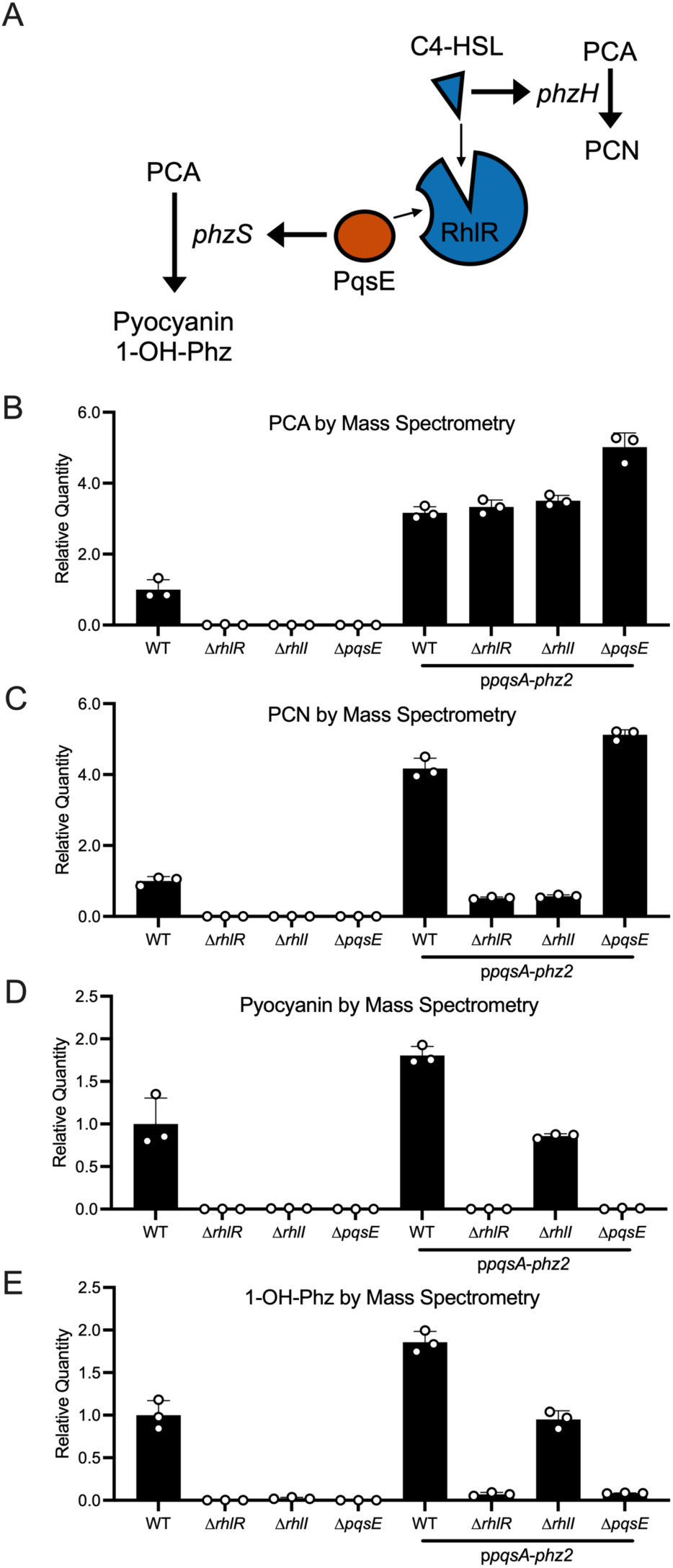
(A) Summary of phenazine regulation by quorum-sensing in *P. aeruginosa*. (B-E) Production of (B) PCA, (C) PCN, (D) Pyocyanin, and (E) 1-OH Phz in the by the designated *P. aeruginosa* strains as measured by mass spectrometry. (B-E) Error bars = standard deviations of biological replicates, *n* = 3.

The compounds identified here inhibit PqsR, and PqsR is required for PqsE production. Thus, we hypothesize that the compounds should inhibit PqsE-RhlR function but not RhlR-C4-HSL function. If so, with respect to target gene expression, the inhibitor compounds should have the same effect as deletion of *pqsE* but not deletion of *rhlI*. To test this supposition, we quantified transcription of *phzH* and *phzS* genes in WT *P. aeruginosa* in the presence and absence of the PqsR inhibitor that we call A8 and compared their levels to their expression levels in the Δ*rhlI*, Δ*pqsE*, and Δ*rhlR* mutants. First, regarding the absence of inhibitor, as previously reported^16,27^ and mentioned above, we find that RhlR-C4-HSL more strongly regulates *phzH* than does PqsE-RhlR as there is lower *phzH* expression in the Δ*rhlI* mutant than in the Δ*pqsE* mutant (Figure S7A). Addition of the A8 inhibitor to the WT drives the *phzH* reduction only to the level displayed by the Δ*pqsE* mutant (Figure S7A). Conversely, here we show that PqsE-RhlR contributes more to *phzS* regulation than does RhlR-C4-HSL (Figure S7B). The Δ*rhlI* strain, which lacks C4-HSL, shows little change in expression of *phzS* compared to that in WT *P. aeruginosa*, demonstrating that RhlR does not require C4-HSL to activate *phzS* transcription. However, when A8 is added to WT *P. aeruginosa*, *phzS* expression is reduced to the level in the Δ*pqsE* strain, demonstrating that indeed, RhlR depends on PqsE to activate *phzS*. Thus, we conclude that compounds that inhibit PqsR interfere with expression of PqsE-RhlR-controlled targets without affecting expression of RhlR-C4-HSL-controlled targets.

In addition to measuring the effects of our compounds on gene expression, ideally, we would quantify how the inhibitors affect production of particular phenazine compounds in the context of RhlR regulation and the individual contributions of PqsE and C4-HSL. However, such an analysis is not straightforward because both *rhlI* and *pqsE* are required for any phenazines to be made^15,16,27^. That is because as described^16,27^ and shown above (Figures 1B and 2B), both *phz1* and *phz2* operon expression rely on RhlI and PqsE. All our attempts to overcome this limitation by constitutively expressing *phz1* or *phz2* were unsuccessful presumably because phenazines are toxic^66,67^, including to *P. aeruginosa*, perhaps explaining why they are under strict quorum-sensing control. To circumvent this issue, we engineered a *P. aeruginosa* strain in which the *pqsA* promoter drives *phz2* operon expression (p*pqsA-phz2*). This strategy places *phz2* operon expression under the exclusive control of PqsR-PQS, enabling density-dependent production of phenazines while eliminating regulation of *phz2* by RhlR. Key for our strategy is that *phzH* and *phzS* are not members of the *phz2* operon, so their regulation by RhlR is maintained. Thus, we can quantify PCA, the product of the *phz2* operon, as well as PCN, pyocyanin, and 1-OH-Phz, which are made from PCA by the action of PhzH and PhzS (Figure 6A). Doing this assessment in the WT, Δ*rhlI*, and Δ*pqsE* strains allows us to measure the contributions of C4-HSL and PqsE to RhlR regulation of *phzH* and *phzS*.

To measure the amounts of particular phenazines produced in the test strains, we analyzed *P. aeruginosa* cell-free culture fluids for the relevant products using liquid chromatography-mass spectrometry. Introduction of p*pqsA-phz2* drove 3-fold increased production of PCA in all strain backgrounds compared that produced by WT *P. aeruginosa* (Figure 6B), showing that the *pqsA* promoter drives higher levels of expression of the *phz2* operon than the native promoter. Deletion of *rhlR* eliminated PCA, PCN, pyocyanin, and 1-OH-Phz production, consistent with a role for RhlR in production of all four phenazines (Figure 6B-E). When we synthetically drove expression of the *phz2* operon, PCN was produced by the Δ*pqsE* strain but not the Δ*rhlI* strain, in agreement with regulation of *phzH* by RhlR-C4-HSL but not by PqsE-RhlR (Figure 6C). By contrast, pyocyanin and 1-OH-Phz production were eliminated in the Δ*pqsE* strain but not the Δ*rhlI* strain, consistent with regulation of *phzS* by PqsE-RhlR (Figure 6D-E). The fact that pyocyanin and 1-OH-Phz production occurred in the Δ*rhlI* strain indicates that PqsE-RhlR can drive production of *phzS* in the absence of C4-HSL. Together, these data show that PqsE-RhlR but not RlhR-C4-HSL is required for *phzS* expression while *phzH* expression requires RhlR-C4-HSL, but not PqsE-RhlR.

## Conclusions

*P. aeruginosa* remains a significant infectious threat, especially among the immune compromised and patients in hospitals. Given that quorum sensing drives pathogenicity via control of toxin production and biofilm formation in *P. aeruginosa*, inhibition of quorum sensing could suppress infectivity, without altering growth, stalling resistance development. Here and earlier^45–49,51–56,59,60,68^, potent PqsR inhibitors possessing a variety of chemical structures have been revealed. The apparent vulnerability of PqsR to small molecule inhibition compared to the resilience of RhlR to interference make it difficult to understand how *P. aeruginosa* profits from RhlR being controlled by both C4-HSL and by PqsE. In this work, we have begun to address this conundrum by disentangling the roles of PqsE and C4-HSL in modulating RhlR transcriptional activation, at least with respect to the generation of different combinations of phenazines. But why make different blends of phenazines? Phenazines harbor shared roles in *P. aeruginosa* biology, for example, as electron acceptors albeit with discrete reactivities and reduction potentials, suggesting that *P. aeruginosa* may reap benefits by producing distinct phenazine blends in different environments^69,70^. Future work can focus on improved development of PqsR inhibitors as potential therapeutics for *P. aeruginosa* and on furthering the understanding of how differential regulation of phenazine production is accomplished, how particular blends of phenazines affect *P. aeruginosa* behaviors and perhaps its survival, and if other traits are differentially affected by PqsR-C4-HSL and PqsE-PqsR.

## Methods

### Strains and growth conditions

Strains used in this study are listed in Table S2. Cells were grown in LB (10 g/L tryptone, 10 g/L NaCl, 5 g/L yeast extract) and M9 (1x M9 salts, 0.5% casamino acids. 0.5% glucose, 1 mM MgSO_4_, 100 µM CaCl_2_) media as specified. Antibiotics were used at the following concentrations: ampicillin (200 µg/mL), kanamycin (100 µg/mL), carbenicillin (400 µg/mL). All strains were grown at 37 °C with aeration. All small molecules presented here are commercially available (Table 1), with the exception of 368A, which was obtained from Macroceutics Inc. and subsequently synthesized by and purchased from WuXi AppTec.

### Reporter construction

Primers used in this study are listed in Table S3. The p*chiC*, p*hcnA*, p*phz1*, p*phz2*, p*rhlA*, p*rpsL*, and p*tac* luciferase reporters were constructed by cloning the ∼500 bp of DNA upstream of each gene of interest upstream of the *lux* genes in pUC18T-mini-Tn7T-Tp^71^ using Gibson assembly. These plasmids were conjugated into *P. aeruginosa* UCBPP-PA14 (designed WT or PA14 in the main text), followed by integration into the chromosome at the att-Tn7 site. Correct integrations were confirmed by colony PCR. Deletions of *P. aeruginosa* genes and introduction of point mutations onto the chromosome were performed as previously reported^30^.

### P. aeruginosa reporter assays

Strains were struck onto LB agar and grown overnight at 37 °C. The following day, individual colonies were isolated and transferred into LB liquid medium followed by incubation at 37 °C. After overnight growth, cultures were diluted 1:1000 into M9 medium. 100 µL of these suspensions were aliquoted into wells of 96 well plates. Plates were incubated for 24 h at 37 °C in a BioSpa Automated Incubator (Agilent). Light production and OD_600_ were quantified from each well at either 15 min or 20 min intervals (Neo2, Agilent). When noted, SPR741 (DC Chemicals) was added at the specified concentration.

### Small molecule inhibitor screen

Initial screening conditions were established at Princeton University and verified at WuXi AppTec. The data shown in Figures 2C, 3B, 3C, and S2 were generated at WuXi AppTec. The screen was also performed at WuXi AppTec. The screening steps are described in the following paragraph. We do note that p*lac-tomato* was inserted into the chromosome of the *P. aeruginosa* screening strain in the intergenic region between *pa14_20500* and *pa14_20510*, as reported previously^16^. p*lac-tomato* was intended to be used as a reporter for cell growth. However, OD_600_ proved to be a sufficiently reliable readout of cell growth and was therefore used in all experiments and is reported in the figures in the Results section. All follow-up work was performed in strains lacking p*lac-tomato*. The presence or absence of p*lac-tomato* did not influence any phenotype measured.

To perform the small molecule inhibitor screen, JSV1378 and JSV1384 were struck onto LB agar and grown overnight at 37 °C. The next day, an isolated colony was selected and grown in LB liquid medium at 37 °C with aeration for 16 h. The cell density was assessed by OD_600_, and the culture OD_600_ was normalized to 0.001 in M9 medium supplemented with 0.5% glucose and 0.5% casamino acids. SPR741 was added at 8 µg/mL. This cell suspension was added to wells of 384 well plates that had been pre-seeded with test compounds, the DMSO negative control, or the compound 368A positive control as described in Results. Test compounds were provided at 40 µM. Plates were briefly subjected to centrifugation to remove air gaps and then incubated at 37 °C. Bioluminescence and OD_600_ were quantified at 0, 12, and 20 h. Outliers were removed based on the quality control algorithm at WuXi AppTec. Plates passed quality control assessment as long as <20% of control wells needed to be removed as outliers.

### Hit confirmation

One thousand small molecules from the initial screen were selected for retesting, following the identical protocol used for the screen. Of these molecules, 751 were assayed for luciferase inhibition using *P. aeruginosa* containing p*tac-lux* instead of p*phz1-lux*. Any compound that inhibited this reporter by at least 50% was eliminated from the pipeline. One hundred and sixty-seven compounds were further analyzed in dilution series to determine IC_50_ values with respect to p*phz1-lux* inhibition. A ten-point, two-fold serial dilution series was made of each test compound in DMSO solvent. The highest tested concentration was 40 µM. These assays were carried out in 384 well plates as described above using both the *P. aeruginosa* p*tac-lux* and p*phz1-lux* reporter strains. 49 candidate compounds exhibiting potent IC_50_ values and of differing structures were purchased for in house analysis.

Molecular weights of compounds purchased for in-house analyses were assessed by LCMS (Table S4). Data for compounds examined in positive mode were acquired on an Agilent 6546 LC/Q-TOF operating in MS (seg) mode. The mobile phase was 50/50 water/acetonitrile with 0.1% formic acid. For the Dual AJ ESI source, the acquisition parameters were as follows: Gas temperature; 275 °C, drying gas flow rate; 12 L/min, nebulizer; 35 psi, sheath gas temperature; 325 °C, sheath gas flow; 12 L/min, and VCap; 3500 V. The instrument was tuned before use. Data for compounds detected in negative mode were acquired on an Agilent 6230 LC/TOF. The mobile phase was 50/50 water/acetonitrile with 4 mM ammonium acetate as the modifier. For the Dual AJ ESI source, the acquisition parameters were as follows: Gas temperature; 275 °C, drying gas; 12 L/min, nebulizer; 25 psi, sheath gas temperature; 325 °C, sheath gas flow; 12 l/min, VCap; 3500 V. The reference masses of *m/z* 112.9855 and 966.0005485 served as internal calibrators throughout runs.

Compound purity data were acquired on a 1290 Infinity II HPLC (Table S4). Compounds were diluted to 10 µM in MeOH and 2 µL were injected onto an Agilent InfinityLab Poroshell 120 Aq-C18 (2.1 x 50 mm, 2.7 µm particle size) column. The flow rate was 0.3 mL/min. The mobile phase was a water-acetonitrile (MeCN) gradient containing 0.1% formic acid. Chromatography was performed as follows: 1 min hold at 5% MeCN, ramp up to 90% MeCN over 11 min, return to 5% MeCN over 1 min. Data were processed using Agilent OpenLAB CDS. Percent purity was calculated by integration of the peaks from the α = 254 nm trace.

### Pyocyanin inhibition assay

WT *P. aeruginosa* and the Δ*pqsE* mutant were struck onto LB agar and grown overnight at 37 °C. Individual colonies were diluted into 5 mL of LB liquid medium. Aliquots were transferred into glass tubes containing test compounds with and without 8 µg/mL SPR741. WT *P. aeruginosa* to which DMSO had been added served as the negative control. The *P. aeruginosa* Δ*pqsE* strain to which DMSO had been added served as the positive control. Compounds were tested at 100 µM. Tubes were shaken for 18 h at 37 °C followed by centrifugation at 15,000 RPM for 5 min. Cell-free culture fluids were collected and assessed for pyocyanin by absorbance at 695 nm. Values were normalized by culture density, using absorbance at OD_600_. Percent inhibition of OD_695_/OD_600_ was normalized to the DMSO control.

### E. coli reporter assays

As previously reported^39^, following overnight growth in LB liquid medium at 37 °C, *E. coli* reporter strains were diluted 1:1000 in LB liquid medium containing 1% arabinose (PqsR) or 0.1% arabinose (LasR and RhlR). The reporter strain for PqsR activity carried pCS26-p*pqsA*-*luxCDABE* and pBAD-*pqsR*. The reporter strain for LasR activity carried pCS26-p*lasB*-*luxCDABE* and pBAD-*lasR*; the reporter strain for RhlR activity carried pCS26-p*rhlA*-*luxCDABE* and pBAD-*rhlR*. Test compounds were added at the specified concentrations. To activate PqsR, PQS was added at 100 nM. To activate LasR, 3O-C12-HSL was added at 5 nM. To activate RhlR, C4-HSL was added at 10 µM. To establish IC_50_ values, 10-point, 10-fold serial dilution series were made starting at 100 µM of each compound. The mixtures were incubated at 37 °C for 24 h (BioSpa, Agilent). Bioluminescence and OD_600_ were assessed every 15 min (Neo2, Agilent). Readings from the 9 h timepoint, representing peak signal production, were used to calculate the IC_50_ values.

Medium conditions used in competition assays are identical to those described above. PQS was assessed in an 11-point, 5-fold dilution series starting at 10 µM. Inhibitors were added as specified, and concentrations were chosen based on their IC_50_ values. To determine EC_50_ values for PQS in the presence of inhibitor, data from the five timepoints flanking the peak of light production from the reporter were averaged. To calculate relative EC_50_ values, each EC_50_ value for PQS at each concentration of inhibitor was normalized to the EC_50_ value determined following addition of the lowest concentration of inhibitor.

### Phenazine quantitation

The designated *P. aeruginosa* strains (Figure 6) were grown as described above for the pyocyanin inhibition assay. Following overnight growth, culture fluids were separated from cells by centrifugation and filtration. As previously reported^72^, cell-free culture fluids were extracted with an equal volume of dichloromethane. The extracts were dried by vacuum evaporation (Genevac HT6 S3i Evaporator) and the samples resuspended in 100% methanol. LCMS data were acquired on an Agilent 1290 Infinity II HPLC connected to an MSD iQ. Dried samples were dissolved into 200 µL methanol, and 2 µL of each sample was injected onto an Agilent InfinityLab Poroshell 120 Aq-C18 (2.1 x 50 mm, 2.7 µm particle size) column. The flow rate was 0.5 mL/min. The mobile phase was a water-MeCN gradient containing 0.1% formic acid. Chromatography was performed as follows: 1 min hold at 5% MeCN, ramp up to 95% MeCN over 5 min, hold at 95% MeCN for 1 min, return to 5% MeCN over 1 min. MS data were acquired in ESI positive mode scan with iQ Auto-Acquire parameters and a capillary voltage of 3500 V. Data were processed using Agilent OpenLAB CDS. Peaks were extracted by *m/z* and compared to a commercial standard of each phenazine. Areas under peaks of interest were quantified.

### Docking hit compounds into the PqsR structure

The PqsR x-ray crystal structure PDB 6Q7U^47^ with its native ligand HHQ was selected for docking studies. The structural data were imported into Maestro [Maestro, version 14.0.136, Schrodinger, LLC, New York, NY, Release 2024-2] from the PDB database. All docking exercises used software modules within the Maestro small molecule suite of applications. The PqsR protein was prepared for docking using the protein preparation workflow with the default settings. Missing side chains were filled in with the Prime module. Water molecules within 5 Å of the ligand were retained. This structure contains a water molecule in the ligand binding pocket that appears hydrogen bonded to the hydroxyl group of Ser-196, the backbone nitrogen of Leu-197, and the carbonyl of the HHQ ligand. This water molecule was essential to dock HHQ reproducibly into its position in the crystal and was maintained for all docking studies. Water molecules more than 5 Å distance from the HHQ ligand were removed. The docking grid was made using Grid Generation by identifying the HHQ ligand within the structure. Docked compounds were confined to the enclosing box center of the centroid of the workspace ligand. Default settings were used in all Receptor Grid Generation tabs except for the Rotatable Groups Setting. Rotatable hydrogen bonding groups from Thr 166, Ser 196, Ser 255, Tyr 258, and Thr 265 located near the ligand binding site were identified and allowed to adopt different orientations using Receptor Grid Generation. The default grid box dimensions of the inner box of 10×10×10 Å and the outer box of 30X30X30 Å were used. All hit compounds were prepared for docking using LigPrep and the OPLS4 force field. Possible ionic states were generated at pH 7.4 ± 1 with Epik and potential tautomers were allowed. The hit compounds were docked into the receptor grid generated from PDB 6Q7U. Compounds were docked in SP mode using the default settings. A maximum of 5 poses per compound were generated and post-docking minimization was performed on 5 poses per ligand. The output was sorted by glide GScore^61^ and the best scoring pose from each compound is shown (Figure 4 and S6). The structure of the compound bound, its best glide GScore, and its pose within the binding site are provided in Table S1. Alignments of previously published PqsR structures were performed in PyMOL [The PyMOL Molecular Graphics System, Version 3.0 Schrödinger, LLC]. All proteins were aligned relative to PDB 6Q7U^47^.

### Real time PCR analysis

Following overnight growth in LB at 37 °C with aeration, *P. aeruginosa* strains were diluted to an OD_600_ of 0.001 and grown to an OD_600_ of 3 at 37 °C with aeration. 500 µL of cells were collected by centrifugation and flash frozen. RNA was extracted using RNAeasy Plus mini columns (Qiagen). DNA was removed, followed by cDNA synthesis, using SuperScript™ IV VILO™ Master Mix with ezDNase™ Enzyme (Invitrogen). RNA was quantified using PerfeCTa SYBR Green FastMix Low ROX (Quanta BioSciences). Outliers with values higher or lower than twice the standard deviation of the mean were removed.

### Data analysis and visualization

Screening data were processed and analyzed in RStudio. Graphs were prepared using Graphpad Prism.

## Supporting Information

Light production over time in *P. aeruginosa* strains carrying 6 additional reporter constructs; percent inhibition of light production from a p*tac-lux* reporter construct in the presence of compounds; inhibition of light production from p*rhlA-lux* and p*lasB-lux* reporters in *E. coli* in the presence of hit compounds; inhibition of pyocyanin production in *P. aeruginosa* by hit compounds in the absence of SPR741; Bemis-Murcko skeleton analyses of hit compounds; docking and ligand competition analyses of hit compounds; quantitative PCR measurements of *phzH* and *phzS* transcription; table summarizing computational results of docking analyses; table summarizing strains used; table summarizing primers used; table summarizing analytical data for purchased compounds (PDF).

## Acknowledgments

We thank members of the Bassler laboratory for helpful discussions. As stated in the text, small molecule screening and initial compound retesting were performed by WuXi AppTec. This work was supported by the Howard Hughes Medical Institute as well as grants from the NIH (R37GM065859) and NSF (MCB-2043238). The content is solely the responsibility of the authors and does not necessarily represent the official views of the National Institutes of Health. The authors declare that they have no competing financial interests.

