## Supporting Information for "Inhibitors of the PqsR Quorum-Sensing Receptor Reveal Differential Roles for PqsE and RhlI in Control of Phenazine Production"

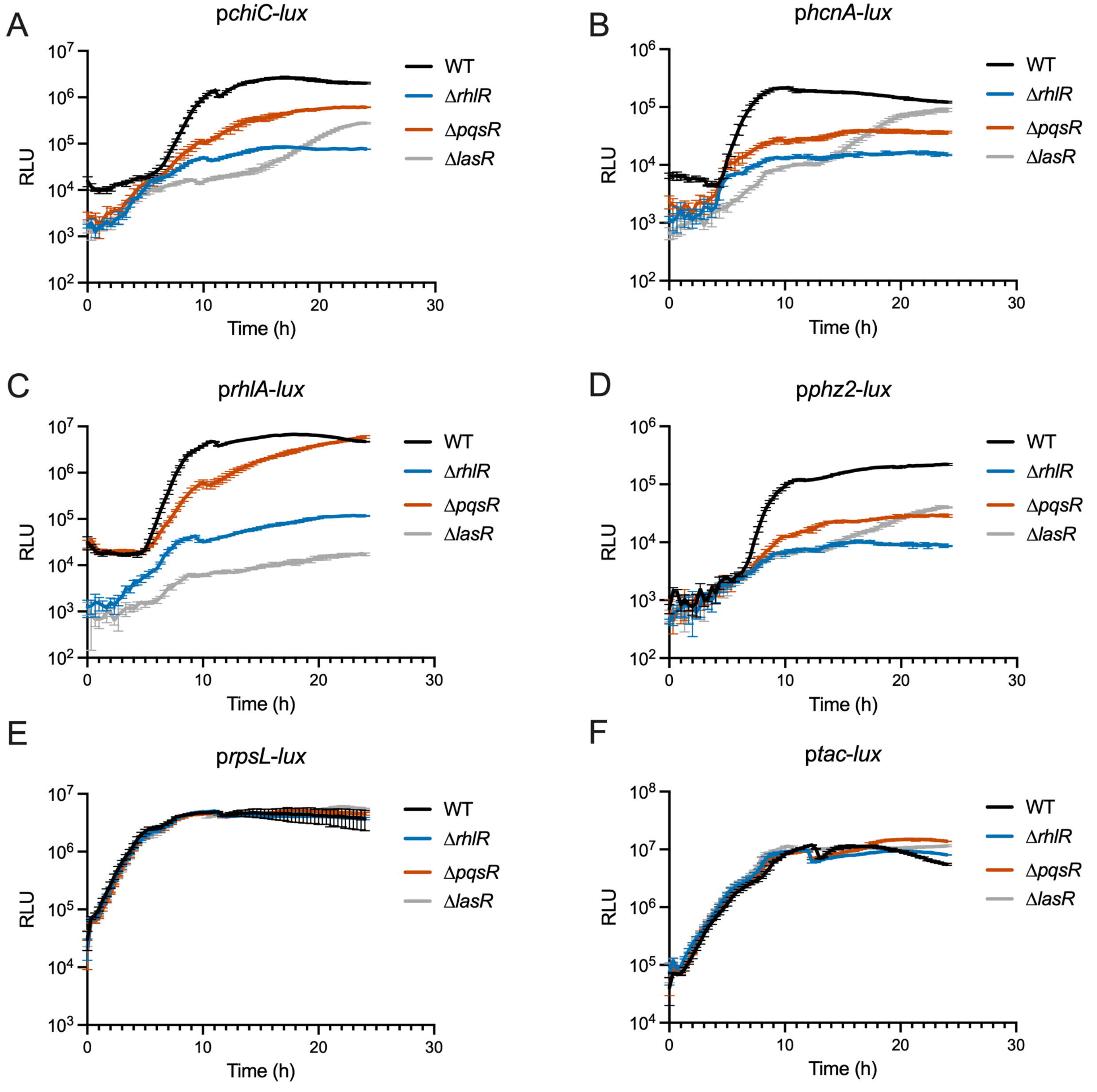


Figure S1. Light production over time in the designated *P. aeruginosa* strains carrying (A) p*chiC*-*lux*, (B) p*hcnA-lux*, (C) p*rhlA-lux*, (D) p*phz2-lux*, (E) p*rpsL-lux*, and (F) p*tac-lux*. RLU as in Figure 1. In all panels, error bars = standard deviations of biological replicates, *n* = 3.

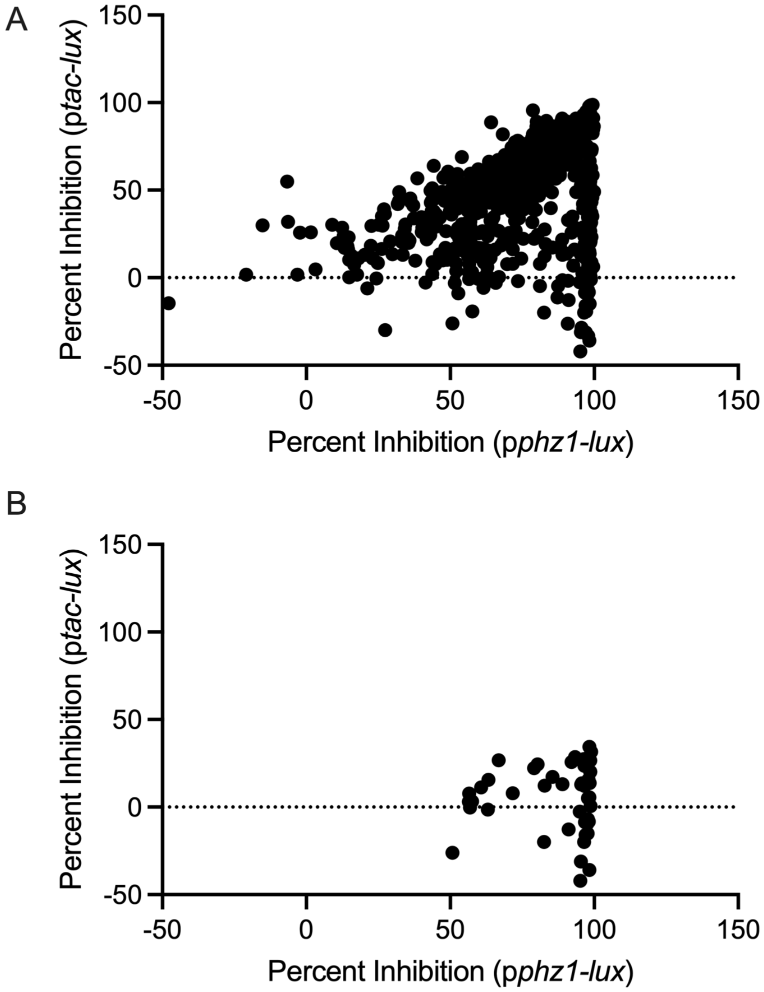


Figure S2. Percent inhibition of light production from the p*tac-lux* reporter versus percent inhibition of light production from the p*phz1-lux* reporter in WT *P. aeruginosa* for (A) all compounds tested in step 3 of the screen and (B) the final compounds from step 5 of the screen.

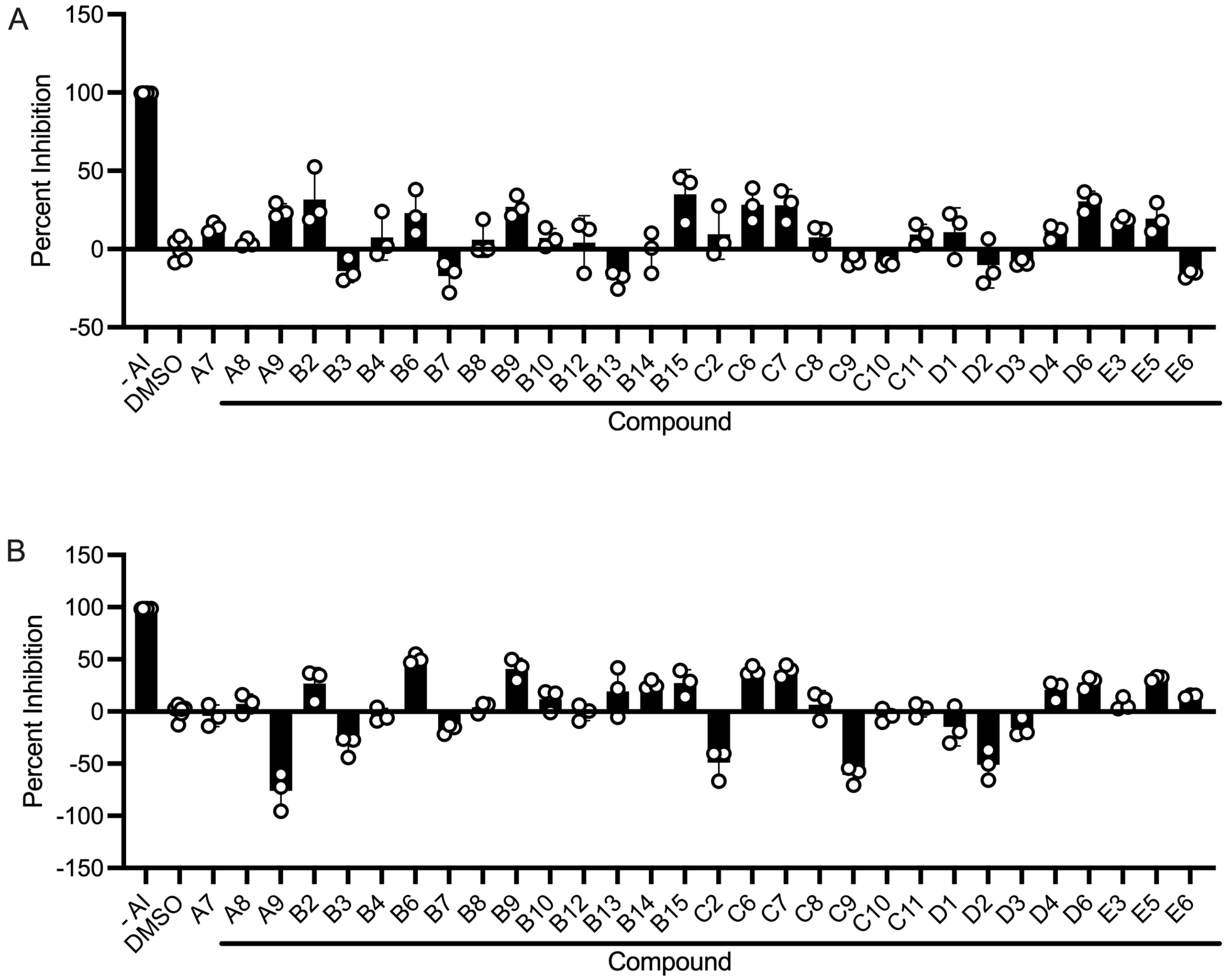


Figure S3. Inhibition of light production from the (A) p*rhlA*-*lux* and (B) p*lasB*-*lux* reporters in *E. coli* by 100 μM of the designated candidate compounds. The left-most bar in each panel shows the results when no C4-HSL (panel A) or no 3O-C12-HSL (panel B) autoinducer was added. For all other bars, either 10 μM C4-HSL (panel A) or 5 nM 3O-C12-HSL (panel B) was present. Error bars = standard deviations of biological replicates, *n* = 3.

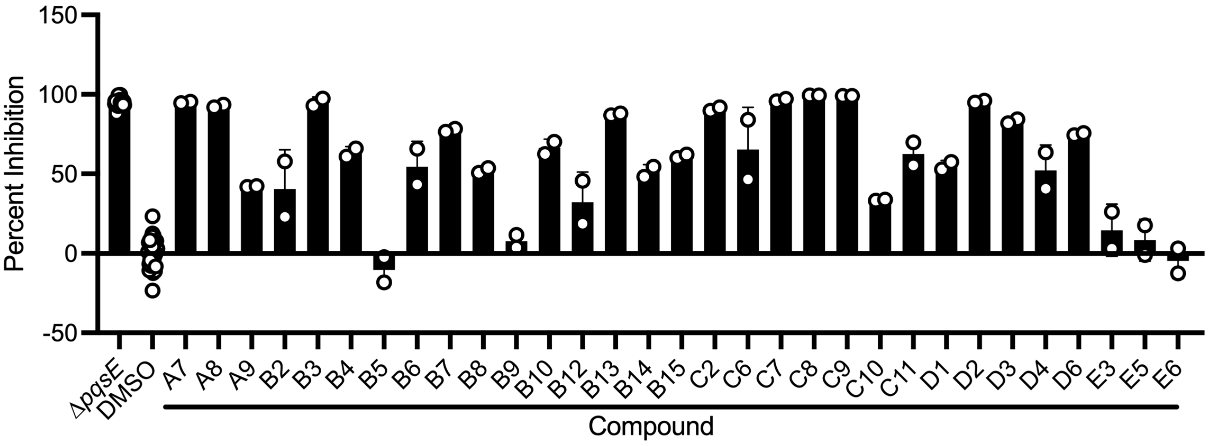


Figure S4. Inhibition of pyocyanin production in WT *P. aeruginosa* by 100 μM of the designated candidate compounds in the absence of SPR741. The left-most bar shows the Δ*pqsE P. aeruginosa* mutant used as the positive control and the second bar shows the result for DMSO as the negative control. Error bars = standard deviations of biological replicates, $n = 3$.

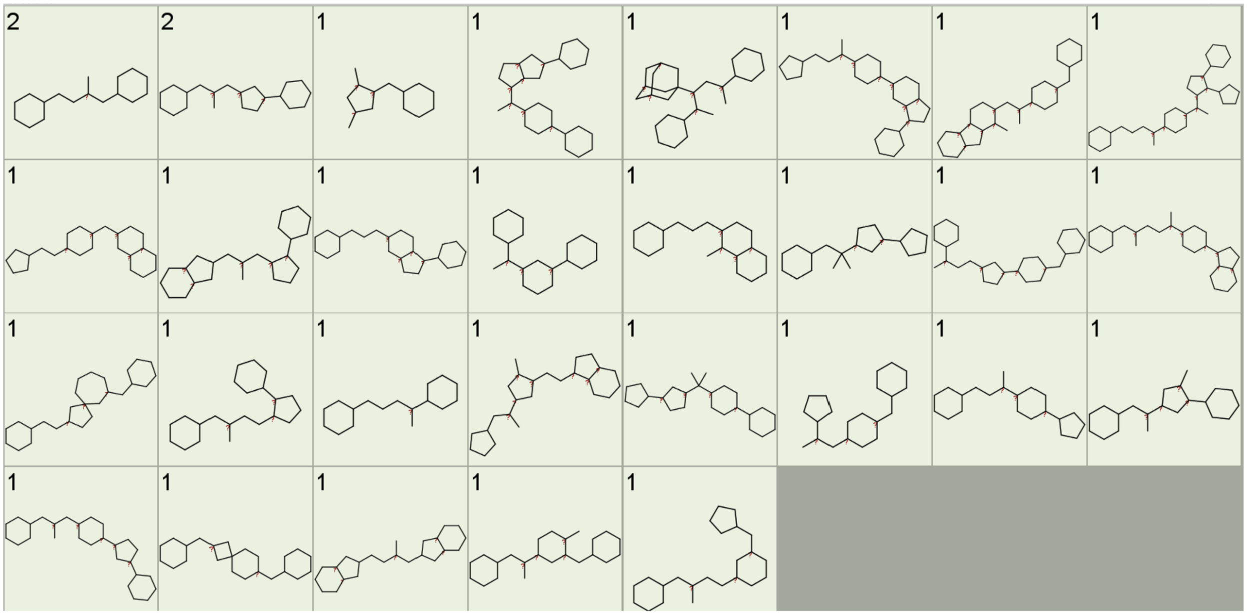


Figure S5. Structural diversity of PqsR inhibitor hits. Bemis-Murcko skeleton analysis of the hits indicated a range of chemical skeletons and a low frequency of duplicate skeletons. Bemis-Murcko skeletons[1] were calculated in DataWarrior (OpenChemLib v06.02.01)[2] and the frequency of each skeleton is shown in the top left corner of each structure box.

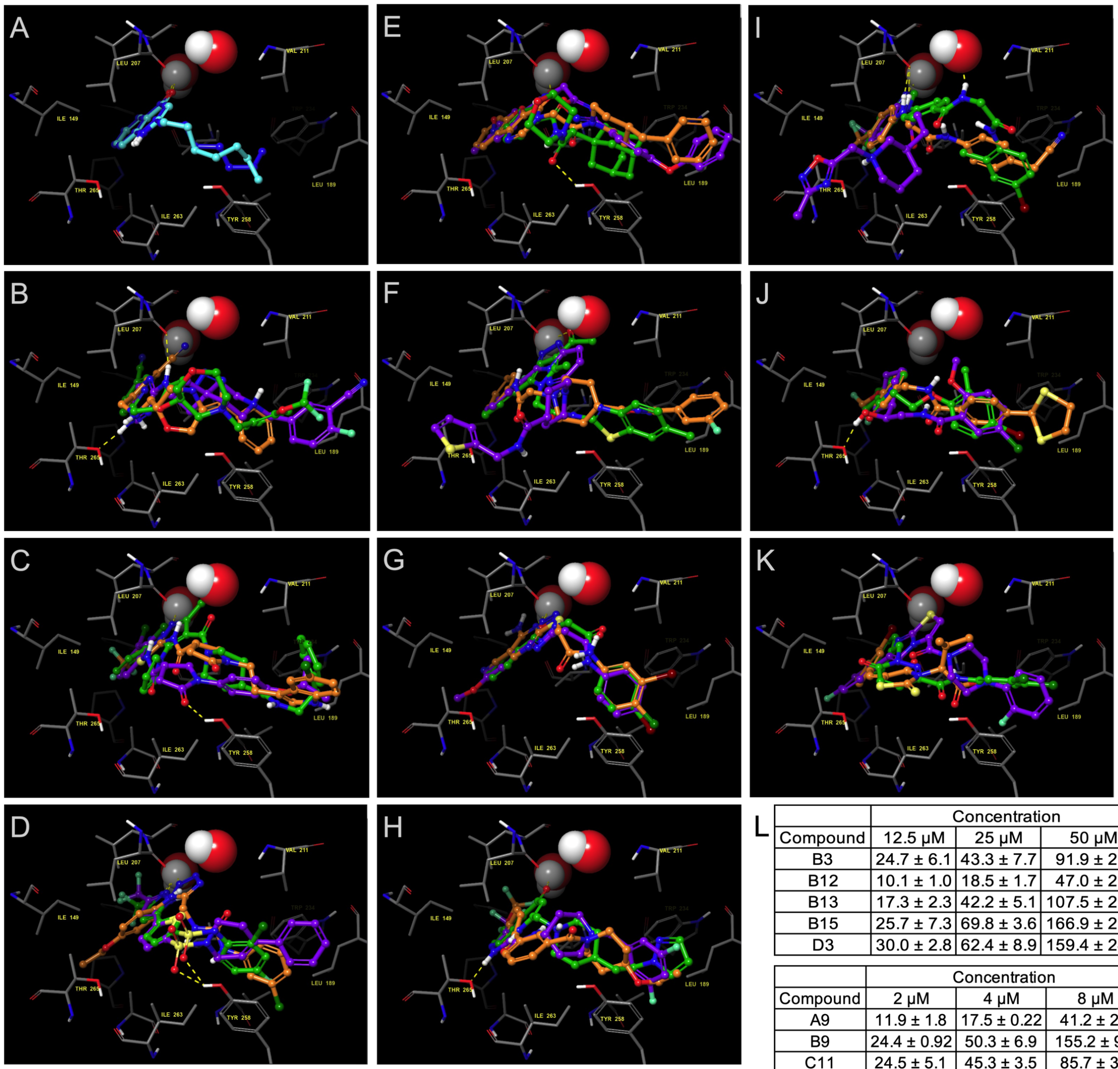


L

| Compound | Concentration 12.5 μM | 25 μM | 50 μM | |
|---|---|---|---|---|
| B3 | 24.7 ± 6.1 | 43.3 ± 7.7 | 91.9 ± 2.7 | EC$_{50}$ (nM) |
| B12 | 10.1 ± 1.0 | 18.5 ± 1.7 | 47.0 ± 2.7 | |
| B13 | 17.3 ± 2.3 | 42.2 ± 5.1 | 107.5 ± 25.0 | |
| B15 | 25.7 ± 7.3 | 69.8 ± 3.6 | 166.9 ± 28.5 | |
| D3 | 30.0 ± 2.8 | 62.4 ± 8.9 | 159.4 ± 24.2 | |

| Compound | Concentration 2 μM | 4 μM | 8 μM | |
|---|---|---|---|---|
| A9 | 11.9 ± 1.8 | 17.5 ± 0.22 | 41.2 ± 2.2 | EC$_{50}$ (nM) |
| B9 | 24.4 ± 0.92 | 50.3 ± 6.9 | 155.2 ± 9.1 | |
| C11 | 24.5 ± 5.1 | 45.3 ± 3.5 | 85.7 ± 3.6 | |
| D2 | 17.7 ± 3.6 | 35.1 ± 3.0 | 109.0 ± 33.0 | |
| D4 | 3.39 ± 0.67 | 9.32 ± 1.6 | 19.1 ± 3.0 | |

| Compound | Concentration 0.5 μM | 1 μM | 2 μM | |
|---|---|---|---|---|
| A7 | 26.1 ± 5.1 | 56.1 ± 12.6 | 91.3 ± 11.0 | EC$_{50}$ (nM) |
| C7 | 19.0 ± 1.6 | 41.1 ± 1.9 | 79.9 ± 7.4 | |
| C8 | 40.5 ± 8.0 | 83.1 ± 3.5 | 185.8 ± 21.3 | |
| C9 | 27.8 ± 2.3 | 77.3 ± 6.4 | 130.8 ± 15.4 | |
| D6 | 7.38 ± 0.82 | 17.0 ± 0.83 | 30.2 ± 10.4 | |

Figure S6. (A) Overlay of the HHQ ligand determined in the PqsR x-ray structure 6Q7U[3] (light blue) and docked into the same protein structure using our docking method (dark blue). (B-K) PqsR inhibitors identified in the current work docked into PqsR x-ray structure 6Q7U. In each panel, water molecules are shown in CPK representation and atoms are colored as follows: oxygen (red), nitrogen (dark blue), hydrogen (white, only hydrogen atoms attached to heteroatoms are shown), sulfur (yellow), chlorine (dark green), fluorine (light green), carbons of PqsR protein (gray). Carbon atoms of docked inhibitor compounds are colored as follows: (B) A7 (purple), A8 (green), A9 (orange); (C) B2 (purple), B3 (green), B4 (orange); (D) B6 (purple), B7 (green), B8 (orange); (E) B9 (purple), B10 (green), B12 (orange); (F) B13 (purple), B14 (green), B15 (orange); (G) C2 (purple), C6 (green), C7 (orange); (H) C8 (purple), C9 (green), C10 (orange); (I) C11 (purple), D1 (green), D2 (orange); (J) D3 (purple), D4 (green), D6 (orange); (K) E3 (purple), E5 (green), E6 (orange). A yellow dashed line indicates a hydrogen bond. (L) $EC_{50}$ values (nM) from competition assays with inhibitors against PQS. These data are presented in Figure 4C-E as values relative to the $EC_{50}$ resulting from addition of the lowest concentration of the given inhibitor.

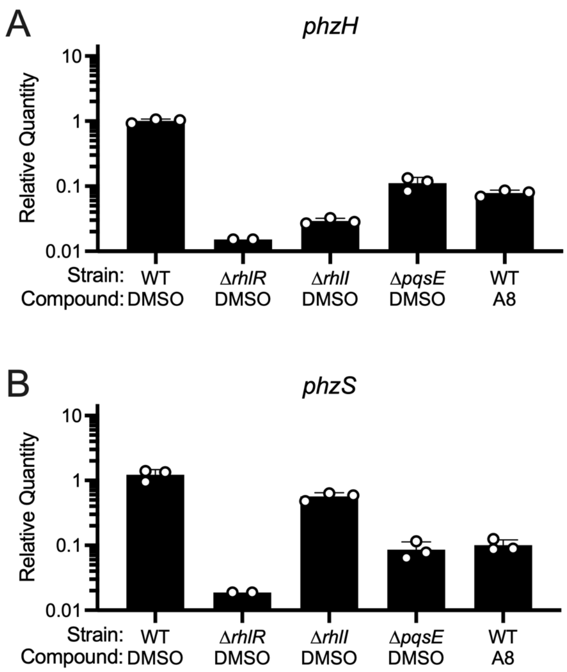


Figure S7. Quantitative PCR measurements of *phzH* (A) and *phzS* (B) in the designated strains in the presence of 1% (v/v) DMSO or the PqsR inhibitor A8 at 100 μM. Error bars = standard deviations of biological replicates, *n* = 3.

Table S1. Docking analysis of PqsR inhibitors

| Compound | Structure | Glide GScore (kcal/mol) | Tot Q (kcal/mol) | State Penalty |
|---|---|---|---|---|
| A7 | 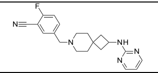 | -6.8 | 1 | 0.01 |
| A8 | 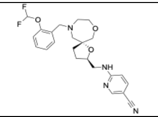 | -7.5 | 1 | 0.30 |
| A9 | 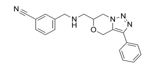 | -7.4 | 1 | 0.33 |
| B2 | 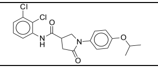 | -7 | 0 | 0.00 |
| B3 | 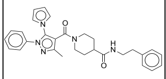 | -6.5 | 0 | 0.00 |
| B4 | 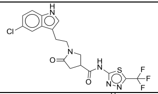 | -7.1 | 0 | 0.37 |
| B6 | 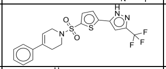 | -6.3 | 0 | 0.30 |
| B7 | 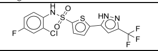 | -6.7 | -1 | 0.41 |

| Compound | Structure | Glide GScore (kcal/mol) | Tot Q (kcal/mol) | State Penalty |
|---|---|---|---|---|
| B8 | 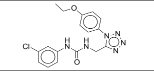 | -6.9 | 0 | 0.00 |
| B9 | 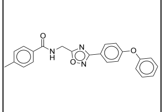 | -7.1 | 0 | 0.00 |
| B10 | 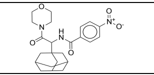 | -5.8 | 0 | 0.00 |
| B12 | 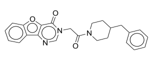 | -6.1 | 0 | 0.00 |
| B13 | 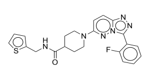 | -6.4 | 0 | 0.00 |
| B14 | 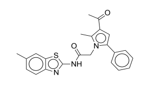 | -6.6 | -1 | 0.81 |
| B15 | 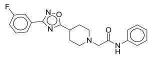 | -7.2 | 1 | 0.47 |
| C2 | 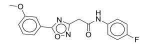 | -7.9 | 0 | 0.00 |

| Compound | Structure | Glide GScore (kcal/mol) | Tot Q (kcal/mol) | State Penalty |
|---|---|---|---|---|
| C6 | 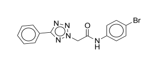 | -7.5 | 0 | 0.00 |
| C7 | 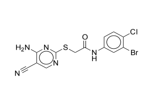 | -6.5 | 0 | 0.00 |
| C8 | 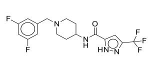 | -8.4 | 0 | 0.49 |
| C9 | 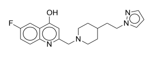 | -8 | 1 | 0.14 |
| C10 | 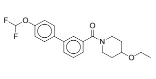 | -5.9 | 0 | 0.00 |
| C11 | 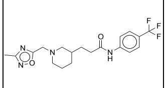 | -7.8 | 1 | 0.00 |
| D1 | 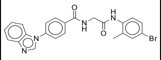 | -7.3 | 0 | 0.01 |

| Compound | Structure | Glide GScore (kcal/mol) | Tot Q (kcal/mol) | State Penalty |
|---|---|---|---|---|
| D2 | 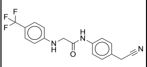 | -7.7 | 0 | 0.00 |
| D3 | 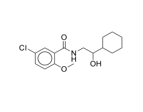 | -6.2 | 0 | 0.00 |
| D4 | 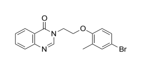 | -7.5 | 0 | 0.00 |
| D6 | 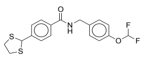 | -6.7 | 0 | 0.00 |
| E3 | 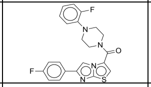 | -7 | 0 | 0.00 |
| E5 | 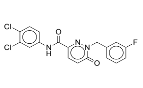 | -6.8 | 0 | 0.00 |
| E6 | 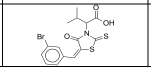 | -5.7 | -1 | 0.00 |

Footnote: Structures of docked PqsR inhibitors with their best docking Glide GScores (kcal/mol), total charge of docked ligand (Tot Q), and state penalty (kcal/mol). Glide GScore is used to predict the binding affinity of a ligand to a target protein[61]. Lower GScores indicate potential for higher binding affinity.

Table S2 - Strain list

| Strain | Genotype | Reference |
|---|---|---|
| JSV1310 | PA14 p*phz1-lux* | This work |
| JSV1563 | PA14 p*phz1-lux* Δ*lasR* | This work |
| JSV1561 | PA14 p*phz1-lux* Δ*lasI* | This work |
| JSV1559 | PA14 p*phz1-lux* Δ*rhlR* | This work |
| JSV1556 | PA14 p*phz1-lux* Δ*rhlI* | This work |
| JSV1534 | PA14 p*phz1-lux* Δ*pqsR* | This work |
| JSV1553 | PA14 p*phz1-lux* Δ*pqsA* | This work |
| JSV1362 | PA14 p*phz1-lux* Δ*pqsE* | This work |
| JSV1558 | PA14 p*phz1-lux* Δ*rhlI* Δ*pqsE* | This work |
| JSV1402 | PA14 p*phz1-lux pqsE*::*pqsE*-D73A | This work |
| JSV1397 | PA14 p*phz1-lux pqsE*::*pqsE*-NI | This work |
| JSV1423 | PA14 p*chiC-lux* | This work |
| JSV1568 | PA14 p*chiC-lux* Δ*rhlR* | This work |
| JSV1538 | PA14 p*chiC-lux* Δ*pqsR* | This work |
| JSV1566 | PA14 p*chiC-lux* Δ*lasR* | This work |
| JSV1269 | PA14 p*hcnA-lux* | This work |
| JSV1570 | PA14 p*hcnA-lux* Δ*rhlR* | This work |
| JSV1536 | PA14 p*hcnA-lux* Δ*pqsR* | This work |
| JSV1591 | PA14 p*hcnA-lux* Δ*lasR* | This work |
| BB-Pa426 | PA14 p*rhlA-lux* | This work |
| JSV1593 | PA14 p*rhlA-lux* Δ*rhlR* | This work |
| JSV1587 | PA14 p*rhlA-lux* Δ*pqsR* | This work |
| JSV1589 | PA14 p*rhlA-lux* Δ*lasR* | This work |
| JSV1269 | PA14 p*phz2-lux* | This work |
| JSV1578 | PA14 p*phz2-lux* Δ*rhlR* | This work |
| JSV1533 | PA14 p*phz2-lux* Δ*pqsR* | This work |
| JSV1586 | PA14 p*phz2-lux* Δ*lasR* | This work |
| JSV1364 | PA14 p*rpsL-lux* | This work |
| JSV1569 | PA14 p*rpsL-lux* Δ*rhlR* | This work |
| JSV1530 | PA14 p*rpsL-lux* Δ*pqsR* | This work |
| JSV1567 | PA14 p*rpsL-lux* Δ*lasR* | This work |
| JSV1390 | PA14 p*tac-lux* | This work |
| JSV1579 | PA14 p*tac-lux* Δ*rhlR* | This work |
| JSV1542 | PA14 p*tac-lux* Δ*pqsR* | This work |
| JSV1583 | PA14 p*tac-lux* Δ*lasR* | This work |
| JSV1378 | PA14 Tn*7*-p*phz1-lux*, *ect6*-p*lac-tomato* | This work |
| JSV1384 | PA14 Tn*7*-p*phz1-lux*, *ect6*-p*lac-tomato* Δ*pqsE* | This work |
| PA14 | WT *P. aeruginosa* | [4] |
| BB-Pa0482 | PA14 Δ*pqsE* | [5] |
| JSV1552 | TOP10 pCS26-p*pqsA-luxCDABE* pBAD-*pqsR* | This work |
| BB-Ec0386 | TOP10 pCS26-p*rhlA-luxCDABE* pBAD-*rhlR* | [6] |
| BB-Ec0382 | TOP10 pCS26-p*lasB-luxCDABE* pBAD-*lasR* | [6] |
| JSV1404 | PA14 Tn*7*-p*tac-lux*, *ect6*-p*lac-tomato* | This work |

| | | |
|---|---|---|
| JSV1629 | PA14 p*pqsA-phz2* | This work |
| JSV1637 | PA14 p*pqsA-phz2* Δ*pqsE* | This work |
| JSV1638 | PA14 p*pqsA-phz2* Δ*rhlI* | This work |
| JSV1643 | PA14 p*pqsA-phz2* Δ*rhlR* | This work |
| JSV1548 | PA14 p*phz1-lux* pUCP18 | This work |
| JSV1549 | PA14 p*phz1-lux* pUCP18-p*lac-pqsE* | This work |
| JSV1544 | PA14 p*phz1-lux* Δ*pqsR* pUCP18 | This work |
| JSV1545 | PA14 p*phz1-lux* Δ*pqsR* pUCP18-p*lac-pqsE* | This work |

Table S3 - Primer list

| Primer name | Primer Sequence | Primer Purpose |
|---|---|---|
| JSV1075 | GCACATCGGCGACGTGCTCTC | Confirm reporter integration |
| JSV1076 | CTGTGCGACTGCTGGAGCTGA | Confirm reporter integration |
| JSV1077 | CACAGCATAACTGGACTGATTTC | Confirm reporter integration |
| JSV1078 | ATTAGCTTACGACGCTACACCC | Confirm reporter integration |
| JSV1059 | ATTTCATTCATTATTAACGGCCAGG TTG | Amplify vector for reporters |
| JSV1060 | GGATCCACTAGTGAGCTCATGC | Amplify vector for reporters |
| JSV1063 | CGATCATGCATGAGCTCACTAGTGG ATCCTTCGATCAGCGTAGTGACG | Amplify p*hcnA* |
| JSV1064 | CTGGCCGTTAATAATGAATGAAATC TTCTTAGTCATTGCCCTTTCATCCGT GAGAG | Amplify p*hcnA* |
| JSV1239 | GAGCTCACTAGTGGATCGTGAACG GCGCG | Amplify p*chiC* |
| JSV1240 | GAATGAAATCTTCTTAGTCATTTAC CGCTCCTCGGATG | Amplify p*chiC* |
| PrhlA-lux F | CGATCATGCATGAGCTCACTAGTGG ATCCGAAGATCTACGCCAACGAAG G | Amplify p*rhlA* |
| PrhlA-lux R | CTGGCCGTTAATAATGAATGAAATC TTCTTAGTCATCTCACACCTCCCAA AAATTTTC | Amplify p*rhlA* |
| JSV1067 | CGATCATGCATGAGCTCACTAGTGG ATCCAAGCGCTCTATTTCGCACTTC TTGCCC | Amplify p*phz1* |
| JSV1068 | CTGGCCGTTAATAATGAATGAAATC TTCTTAGTCATGCGCCGCCTCCGAG AGGGCTC | Amplify p*phz1* |
| JSV1069 | CGATCATGCATGAGCTCACTAGTGG ATCCGCAAGCTCAACTCCAGCAAC A | Amplify p*phz2* |

| | | |
|---|---|---|
| JSV1070 | CTGGCCGTTAATAATGAATGAAATCTTCTTAGTCATGGTGCGAATCTCCGCCAGTTC | Amplify p*phz2* |
| JSV1166 | CGATCATGCATGAGCTCACTAGTGGATCAATCAACGACAAGCACATCG | Amplify p*rpsL* |
| JSV1167 | CTGGCCGTTAATAATGAATGAAATCTTCTTAGTCATCTATAGCTCCACTGATTGTCTTACG | Amplify *rpsL* promoter |
| JSV1216 | CATTCCACTTACAATTAGGCAAAGGATATGTAGGCTGGAGCTGCTTC | Amplify p*tac* promoter + luciferase |
| JSV1217 | GCATGAGCTCACTAGTGGATCCTGCACCAATGCTTCTGGC | Amplify p*tac* promoter + luciferase |
| JSV1215 | GAAGCAGCTCCAGCCTACATATCCTTTGCCTAATTGTAAGTGGAATG | Amplify vector for p*tac* reporter |
| JSV1218 | GCCAGAAGCATTGGTGCAGGATCCACTAGTGAGCTCATGC | Amplify vector for p*tac* reporter |
| JSV1501 | GGGAACGTTCTGTCATGCGAGAGTACCAACG | Replace p*phz2* with p*pqsA* |
| JSV1502 | TGGTACTCTCGCATGACAGAACGTTCCCTCTTC | Replace p*phz2* with p*pqsA* |
| JSV1503 | GGGGATCCTCTAGAGAATAATCATTAATCAGGTGGGAATG | Replace p*phz2* with p*pqsA* |
| JSV1504 | CGTTTGCGCCGGTTCGCCCTTCTTGCTTGG | Replace p*phz2* with p*pqsA* |
| JSV1506 | AAGCAAGAAGGGCGAACCGGCGCAAACG | Replace p*phz2* with p*pqsA* |
| JSV1508 | ATTAATGATTATTCTCTAGAGGATCCCCGGG | Replace p*phz2* with p*pqsA* |
| JSV1509 | TGTAAAGCAAGCTTGTTTCTTCTGGGAGGAAGC | Replace p*phz2* with p*pqsA* |
| JSV1509 | CTCCCAGAAGAAACAAGCTTGCTTTACATTTATGCTTC | Replace p*phz2* with p*pqsA* |
| JSV1195 | CAAAACTGCCCGCAACGT | *rpsL* quantitative PCR |
| JSV1196 | TTTCGGCGTGGTGGTGTAT | *rpsL* quantitative PCR |
| JSV1551 | GATCTTCGCCATGACCGATAC | *phzH* quantitative PCR |
| JSV1557 | GTGGGAAAGCGATAGGACATC | *phzH* quantitative PCR |
| JSV1443 | CGTCGGCATCAATATCCAGC | *phzS* quantitative PCR |
| JSV1444 | CGTCGGCATCAATATCCAGC | *phzS* quantitative PCR |

Table S4. Analytical data for purchased compounds

| Compound | Ion Type | Expected m/z | Observed m/z | ppm Error | Percent Purity |
|---|---|---|---|---|---|
| A07 | [M+H] | 352.1932 | 352.1959 | -7.72 | >80% |
| A08 | [M+H] | 445.2046 | 445.2073 | -6.11 | >95% |
| A09 | [M+H] | 346.1663 | 346.1679 | -4.68 | >95% |
| B10 | [M+H] | 428.2180 | 428.2199 | -4.48 | >95% |
| B12 | [M+H] | 402.1812 | 402.1832 | -5.02 | >95% |
| B13 | [M+H] | 437.1555 | 437.1586 | -7.14 | >95% |
| B14 | [M-H] | 402.1281 | 402.1289 | -1.92 | >95% |
| B15 | [M+H] | 381.1722 | 381.1742 | -5.30 | >95% |
| B02 | [M-H] | 405.0778 | 405.0790 | -2.89 | >95% |
| B03 | [M+H] | 482.2551 | 482.2572 | -4.40 | >95% |
| B04 | [M+H] | 458.0660 | 458.0679 | -4.19 | >95% |
| B06 | [M+H] | 440.0709 | 440.0700 | 2.00 | >95% |
| B07 | [M-H] | 423.9610 | 423.9626 | -3.71 | >60% |
| B09 | [M+H] | 386.1499 | 386.1480 | 4.87 | >95% |
| C10 | [M+H] | 376.1719 | 376.1739 | -5.37 | >95% |
| C11 | [M+H] | 397.1846 | 397.1866 | -5.09 | >95% |
| C02 | [M+H] | 328.1092 | 328.1111 | -5.85 | >95% |
| C06 | [M-H] | 356.0152 | 356.0161 | -2.45 | >95% |
| C07 | [M+H] | 397.9473 | 397.9487 | -3.57 | >95% |
| C08 | [M+H] | 389.1396 | 389.1420 | -6.22 | >95% |
| C09 | [M+H] | 355.1929 | 355.1945 | -4.56 | >95% |
| D01 | [M+H] | 463.0764 | 463.0784 | -4.36 | >95% |
| D02 | [M+H] | 334.1162 | 334.1179 | -5.15 | >95% |
| D03 | [M+H] | 312.1361 | 312.1369 | -2.63 | >95% |
| D04 | [M+H] | 359.0390 | 359.0392 | -0.61 | >95% |
| D06 | [M+H] | 382.0742 | 382.0758 | -4.24 | >95% |
| E03 | [M+H] | 425.1242 | 425.1258 | -3.81 | >95% |
| E05 | [M+H] | 392.0364 | 392.0385 | -5.41 | >95% |
| E06 | [M-H] | 397.9525 | 397.9535 | -2.44 | >95% |
